# Bactofilins are essential spatial organizers of peptidoglycan insertion in the Lyme disease spirochete *Borrelia burgdorferi*

**DOI:** 10.1101/2025.04.09.647816

**Authors:** Christopher B. Zinck, Valentina Carracoi, Zachary A. Kloos, Jenny Wachter, Cindi L. Schwartz, Philip E. Stewart, Christine Jacobs-Wagner, Patricia A. Rosa, Constantin N. Takacs

## Abstract

The Lyme disease spirochete *Borrelia burgdorferi* has a distinctive pattern of growth. Newly-born cells elongate by primarily inserting peptidoglycan at mid-cell, while in longer cells, additional insertion sites form at the one-quarter and three-quarter positions along the cell length. It is not known how peptidoglycan insertion is concentrated at these locations in *B. burgdorferi.* In other bacteria, multi-protein complexes are known to synthesize new peptidoglycan and are often organized by cytoskeletal proteins. We show here that *B. burgdorferi*’s zonal concentration of peptidoglycan insertion requires BB0538 (BbbA) and BB0245 (BbbB), two members of the bactofilin class of cytoskeletal proteins. Bactofilin depletion redistributes peptidoglycan insertion along the cell length. Prolonged bactofilin depletion arrested growth in culture and induced extensive cell blebbing, indicating that *B. burgdorferi* bactofilins are essential for viability. Fluorescent protein fusions of BbbA and BbbB localized to areas of peptidoglycan insertion, with BbbB accumulation preceding peptidoglycan insertion at these sites. Similar to peptidoglycan insertion, BbbB localization was disrupted upon depletion of BbbA. Our results show that BbbB relies on BbbA for its localization, and that together, BbbA and BbbB direct the spatial patterning of new peptidoglycan insertion in *B. burgdorferi*.

**IMPORTANCE:** The spirochetal bacterium *Borrelia burgdorferi* causes Lyme disease, the most prevalent vector-borne infection in North America and Europe. Cellular replication, which requires growth and division of the peptidoglycan cell wall, facilitates *B. burgdorferi* transmission to, and dissemination within, new hosts. Cellular replication is therefore essential for pathogenesis. Bactofilins regulate peptidoglycan-related processes in several bacteria. However, these functions are typically non-essential for cellular replication, as bactofilin-encoding genes can be readily deleted in multiple bacterial species. In contrast, we show that the *B. burgdorferi* bactofilins BbbA and BbbB are essential for cellular viability and direct zonal peptidoglycan insertion. Our findings broaden the spectrum of known bactofilin functions and advance our understanding of how peptidoglycan insertion is regulated in this unusual, medically important spirochete bacterium.

## INTRODUCTION

*Borrelia burgdorferi* and related spirochetal bacteria are zoonotic pathogens vectored by ixodid ticks (1, 2). These bacteria cause Lyme disease, also known as Lyme borreliosis, the most prevalent vector-borne infectious disease in temperate climates of the Northern Hemisphere (3–6). Larval ticks typically acquire *B. burgdorferi* when they feed on an infected vertebrate animal and then maintain the spirochetes through both the larva-to-nymph and nymph-to-adult molts (7, 8). Ticks transmit *B. burgdorferi* to vertebrate hosts during post-acquisition feedings (9, 10). In the host, *B. burgdorferi* disseminates from the skin to multiple tissues, including distant skin sites, the heart, joints, and the meningeal lining of the central nervous system (11–16). Lyme borreliosis can present with generalized or localized symptoms including fever, malaise, skin rash, heart block and arrhythmias, meningitis and Bell’s palsy, as well as arthritis (3, 17–21). While antibiotic treatment is a generally effective therapy (22–24), post-infectious symptoms can persist in 10-20% of patients and can be debilitating (25–27).

*B. burgdorferi* cellular replication is required for the spirochete’s colonization of and persistence in its tick vectors and vertebrate hosts. The bacterium replicates in the tick midgut during tick feeding and digestion of the blood meal (7, 8, 28), and likely even in post-molt unfed nymphs (29, 30). In the host, *B. burgdorferi* replication underpins its effective dissemination from the bite site, colonization of distal tissues, and long-term antigenic variation-mediated evasion of the host adaptive immune response (14, 31–36). Bacterial replication requires regulated growth and division of the peptidoglycan sacculus, a mesh-like macromolecular polymer made of glycan chains cross-linked by short stem peptides (37–39). Peptidoglycan lies outside the cytoplasmic membrane, opposes the turgor pressure of the cytoplasm, and helps maintain cellular shape and integrity (38). In spirochetes, both peptidoglycan and motility-generating flagella are found within the periplasmic space delimited by the inner and outer membranes (40–50).

Previous studies in several model bacteria have identified two widely conserved protein complexes that carry out peptidoglycan biosynthesis and attendant cell growth. The ubiquitous bacterial divisome complex synthesizes a cell wall septum that separates the two offspring cells during cytokinesis (37, 51, 52). Additionally, in many rod-shaped bacteria, the elongasome contributes to cell growth by inserting new peptidoglycan along the sidewall (37, 38, 53). Alternatively, growth can occur through elongasome-independent, polar synthesis of new peptidoglycan (54–60). Both the divisome and the elongasome are organized by bacterial cytoskeletal elements (68) and contain transglycosylases (e.g., FtsW or RodA, respectively), transpeptidases (e.g., FtsI/PBP3 or PBP2), and regulatory, non-enzymatic components (e.g., FtsN or MreCD) (37, 61, 62). Tubulin homolog FtsZ governs assembly of the divisome (52), while actin homolog MreB orients the activity of the elongasome perpendicular to the long axis of rod-shaped bacteria (63–65). Additionally, bacterial intermediate filament proteins can modulate cell growth (66), for example by associating with the plasma membrane and exerting a mechanical force that directs preferential peptidoglycan insertion on the opposite sidewall (67–69).

Another class of cytoskeletal proteins broadly represented among bacteria, archaea, and some eukaryotes are the bactofilins (70–72). They share a triangular, parallel beta barrel structure formed by the conserved domain of unknown function (DUF) 583 (70, 72, 73). This structure assembles head-to-head and tail-to-tail in the absence of any cofactor to yield nonpolar protofilaments approximately 3 nm-wide (70). These protofilaments can, in turn, interact laterally to form bundles of varying thickness or two-dimensional crystalline lattices (70, 74, 75). The DUF583 bactofilin domain is flanked by N- and C-terminal tails of varying length that mediate protein-protein interactions and membrane association (70, 74, 76, 77).

Bactofilins serve as scaffolds for diverse cellular processes, including motility, biofilm formation, establishment and maintenance of cell polarity, and chromosome segregation (72, 78–81). They have also been suggested or demonstrated to participate in cell growth and morphogenesis processes such as cell division (72) and cell size regulation (82), respectively. In other bacteria, the bactofilins control the growth and shape of cell appendages such as stalks or hyphae (72, 76, 83), or a peculiar mode of cell growth that comprises of budding of a new cell from the free end of a stalk (84). Disruption of bactofilin function has also been shown to induce kinks in otherwise straight rod-shaped cells (85, 86), modify the radius of cell curvature (84–87), or change a cell’s helical pitch (88). These findings established bactofilins as key modulators of the bacterial rod shape.

The mechanisms underpinning bactofilin function remain poorly understood, but several patterns have begun to emerge. Membrane association and filament formation are essential bactofilin properties (70). Additionally, evidence exists of a phylogenetically conserved, functional interaction between some bactofilins and M23 endopeptidases, which are peptidoglycan hydrolases (76, 84, 87, 89). In *Hyphomonas neptunium* and *Rhodospirillum rubrum,* bactofilin BacA recruits an M23 endopeptidase, LmdC, to modify cell shape (84). In *Helicobacter pylori*, the bactofilin CcmA modifies the extent of peptidoglycan crosslinking, possibly through indirect regulation of the stability of the M23 peptidase Csd1 (87, 90, 91). In contrast, bactofilin BacA of *Caulobacter crescentus* interacts with a peptidoglycan synthase, PbpC (72). This finding raises the possibility that bactofilins may regulate peptidoglycan synthases in the other species in which they are known influence peptidoglycan insertion (86, 92).

*B. burgdorferi* cells have a distinct pattern of growth. While peptidoglycan insertion occurs at low levels along most of the cell body, except the inert poles, notably higher insertion occurs at mid-cell (93, 94). As the cells grow, new zones of localized insertion develop at the one-quarter and three-quarter cell positions and the mid-cell zone becomes a division site (93). This pattern of localized cell wall insertion establishes, in one generation, the primary growth zones and eventual division sites of the next generation of cells (93). A CRISPR interference-based gene expression knockdown approach implicated the elongasome in peptidoglycan synthesis at these locations, as depletion of MreB and RodA caused localized bulging (a typical elongasome inhibition phenotype) at the one-quarter, mid-cell, and three-quarter positions (95). Given the paucity of knowledge concerning mechanisms of cell growth in *B. burgdorferi* and the finding that spirochete *L. biflexa* employs bactofilin LbbD in regulating cell helical pitch, cell wall strength, and motility (88), we investigated the functions of two *B. burgdorferi* bactofilins using genetic and imaging approaches.

## RESULTS

### *B. burgdorferi* encodes three bactofilins

Several studies have reported that spirochetes encode bactofilins (70, 72, 84, 88). Leptospirae encode five paralogs, LbbA through LbbE, of which LbbD has been characterized (88). Using Basic Local Alignment Search Tool (BLAST) searches targeting the spirochete phylum, we identified 962 unique spirochete bactofilin sequences, which we used to generate a phylogenetic tree (Fig. 1). Based on their similarity to the previously described *L. biflexa* bactofilins LbbA through LbbE (88), we assigned the spirochete bactofilins to five corresponding clusters, designated A through E (Fig. 1). We found that only the Leptospiraceae encode members of all five clusters, while the Brachyspiraceae encode only cluster D and E bactofilins, and the Spirochaetaceae, Treponemataceae, Breznakiellaceae, and Borreliaceae encode only cluster A, B, and C bactofilins (Fig. 1). The five homologs present in Leptospiraceae are likely ancestral, while the other spirochete families likely lost bactofilin genes during evolution. Within the Borreliaceae, the bactofilin loci were well-conserved among the 28 Lyme disease spirochete genomes we visualized (Fig. S1A) using the BorreliaBase online resource (96). In the *B. burgdorferi* type strain B31, the three bactofilins, chromosomally encoded by genes *bb0538*, *bb0245*, and *bb0231* (Fig. S1A), were expressed in culture (Figs. S1B,C). We propose renaming these genes as *<u>B</u>. <u>b</u>urgdorferi* <u>b</u>actofilins <u>A</u>, <u>B</u>, and <u>C</u> (*bbbA*, *bbbB*, and *bbbC*, respectively) (Figs. 1 and S1A), and will refer to them as such throughout the remainder of the manuscript.

**Figure 1.**
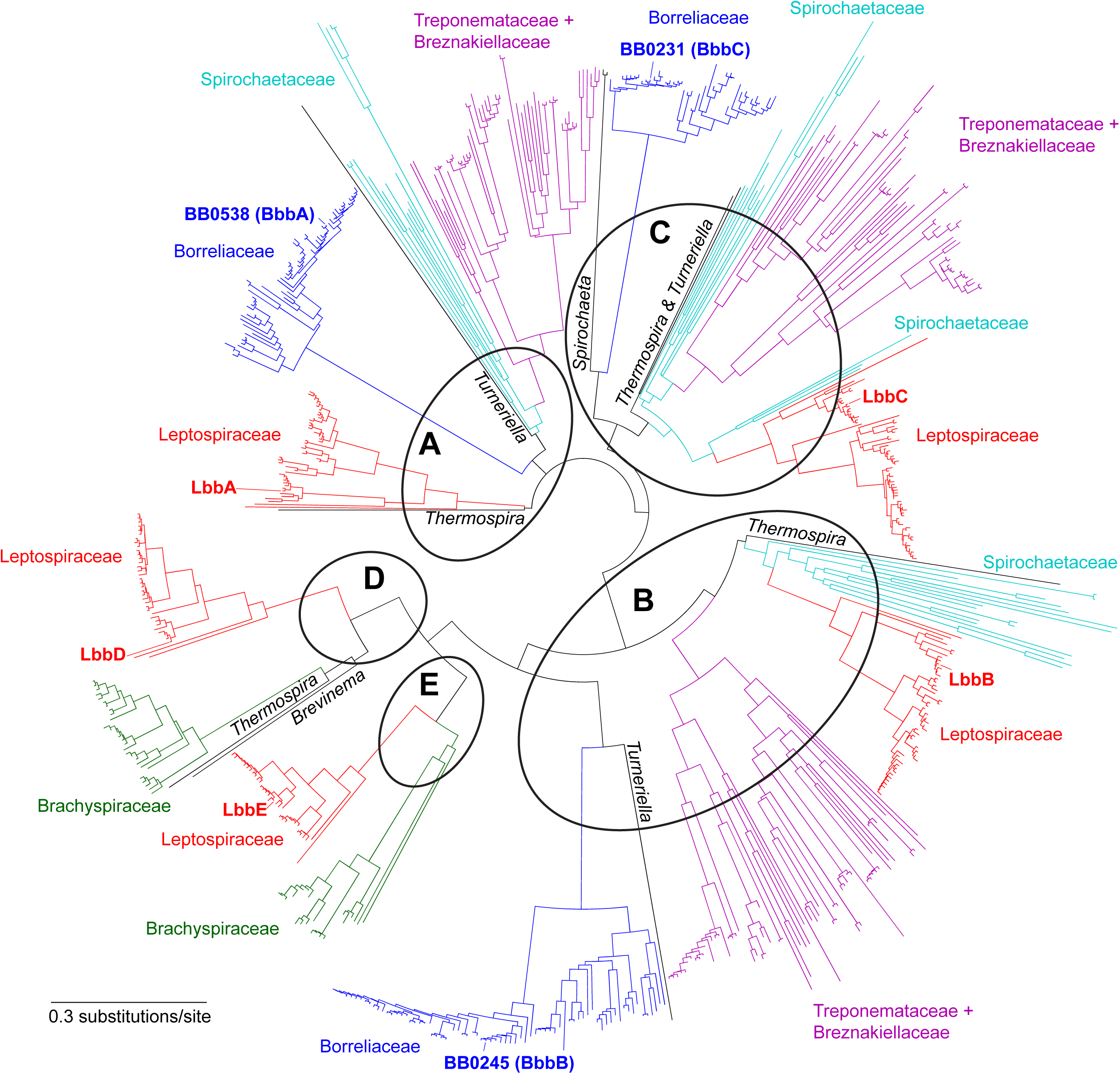
Phylogenetic tree of bactofilin sequences encoded by spirochetes. The tree was generated using 692 non-redundant sequences. Colored branches indicate bactofilin clades belonging to the listed families. Black branches and accompanying genera names indicate sequences that clustered away from the main clades. Circles indicate the grouping of the bactofilin sequences in relation to the five *Leptospira biflexa* proteins LbbA through LbbE, which are indicated on the tree. The three bactofilins encoded by *B. burgdorferi* strain B31, BbbA through BbbC, are also indicated.

Alignment of 174 unique bactofilin sequences belonging to both Lyme disease and relapsing fever *Borrelia* species revealed a central domain (Fig. 2A, gray bar) with high amino-acid conservation (Fig. 2B). AlphaFold3 (97) structural modeling of BbbA, BbbB, and BbbC from *B. burgdorferi* strain B31 predicted that the central domain of each protein folds into a triangular beta-barrel (Fig. 2C), which is the structural hallmark of bactofilins (73, 75, 98). This central, conserved beta-barrel domain was flanked by N- and C-terminal tails (Fig. 2A) that were more variable in sequence and length (Fig. 2B) and predicted to be less structured (Fig. 2C). A subset of the BbbA sequences (24 out of 60) had an N-terminal tail extension of 14 amino-acids (Fig. 2B, star). These extended BbbA sequences contained a conserved second initiation codon at position 15, which aligned with the initiation codon of the shorter BbbA sequences (Fig. S2A, arrowhead). We could not identify a canonical ribosomal binding site (RBS) within thirty base pairs upstream of the annotated initiation codon of gene *bbbA* (*bb0538*) in the genome of the *B. burgdorferi* B31 strain (Fig. S2B). While the existence of leaderless transcripts in *B. burgdorferi* implies translation can occur in the absence of a canonical RBS (99), we found that the second initiation codon of *bbbA* (Fig. S2B, arrowhead) is preceded by a likely RBS (Fig. S2B, blue letters) with a nucleotide sequence (5’-AGAGG-3’) similar to that of the consensus *B. burgdorferi* RBS, 5’-AGGAG-3’ (99). Therefore, for BbbA we performed all analyses and built all genetic constructs assuming translation starts at the M15 position of the annotated sequence (Fig. S2).

**Figure 2.**
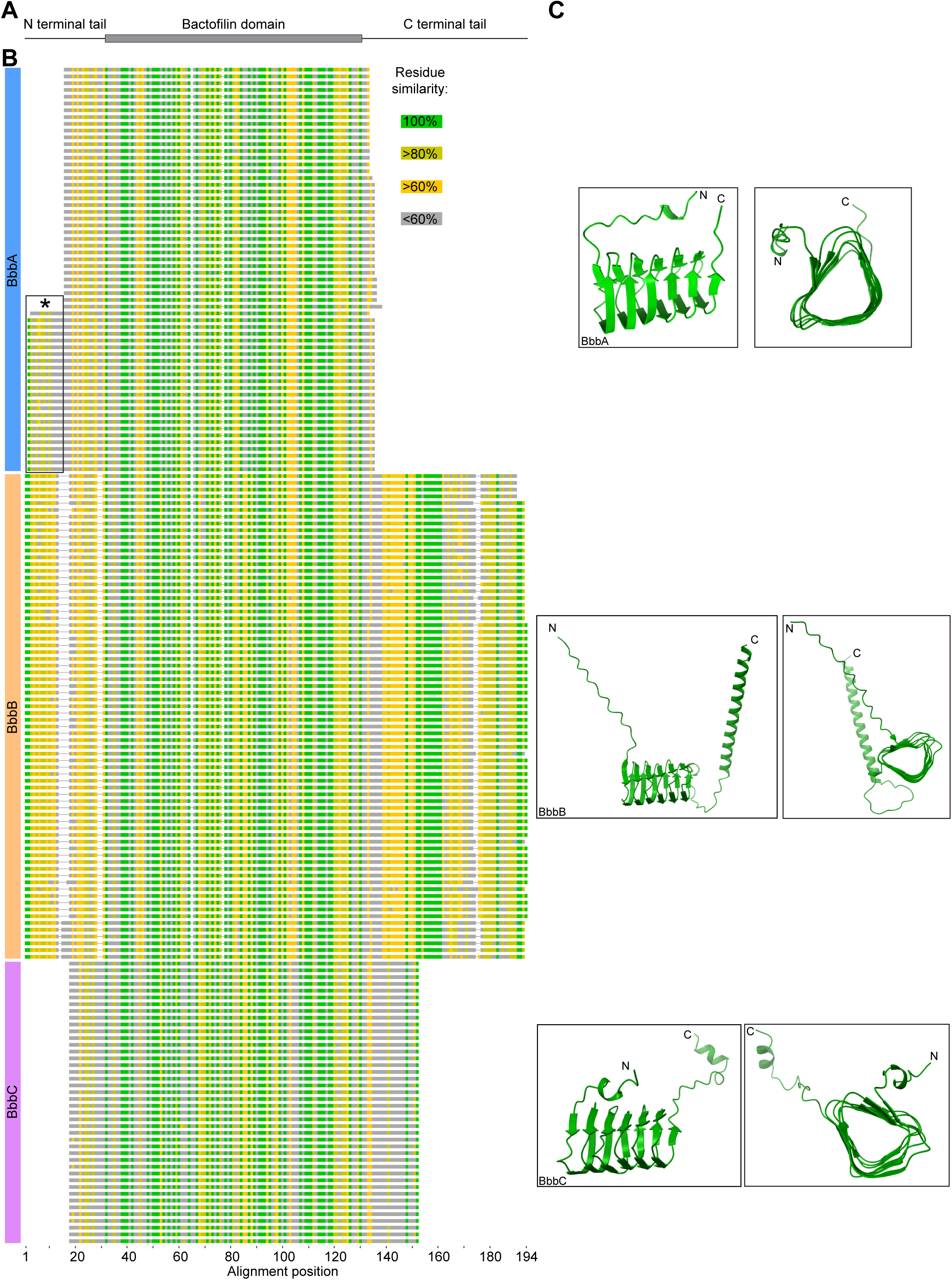
Computational analyses of Borreliaceae bactofilin sequences. **A.** Schematic depiction of domain organization of *B. burgdorferi* bactofilins, matching the sequence alignment shown in panel B. **B.** Sequence alignment of 174 bactofilin sequences encoded by Borreliaceae. The three clades, BbbA, BbbB, and BbbC, are indicated at the left. A subset of BbbA sequences have an extended N-terminal tail, marked by a star (*) sign. Color map for residue similarity levels are at the right. **C.** Alphafold3 structural prediction models for *B. burgdorferi* strain B31 bactofilins BbbA with shortened N-terminal tail (top), BbbB (middle), and BbbC (bottom). Amino (N) and carboxy (C) termini of the polypeptide chains are indicated. Shown are views from the side of the bactofilin domain (left) and along the central axis of the bactofilin domain (right).

### BbbB is required for *B. burgdorferi* growth in culture and maintenance of normal cell morphology

To investigate bactofilin BbbB, we created depletion strain VCbb08 by placing the native *bbbB* gene under the control of the isopropyl β-D-1-thiogalactopyranoside (IPTG)-inducible promoter P*_flac_* (100) (Fig. S3). This strain also constitutively expresses LacI (Fig. S3) (101), which binds the *lacO* sequence within the P*_flac_* promoter and downregulates (but not necessarily abolishes) gene expression. LacI-mediated transcriptional repression is relieved by addition of IPTG to the growth medium (100, 101).

In the presence of 2 mM or 100 μM IPTG, cultures of strain VCbb08 displayed growth kinetics similar to those of the uninduced parental strain B31-68-LS (Fig. 3A). Cells of VCbb08 also had typical spirochetal cell morphology when grown with 100 µM IPTG (Fig. 3B). In contrast, incubation of VCbb08 in culture medium supplemented with 5 µM IPTG or lacking IPTG impaired spirochete growth (Fig. 3A). Additionally, two days of incubation without IPTG caused severe membrane blebbing at varied locations along the cell length (Figs. 3B,C). Continued incubation without IPTG eventually led to a recovery of spirochete growth (Fig. 3A), suggesting that compensatory mutations had arisen in a subpopulation of cells and that these spirochetes had become predominant in the experimental cultures. Mutations that alleviate LacI-mediated repression have been described previously when this system has been used for conditional expression of other essential genes in *B. burgdorferi* (102).

**Figure 3.**
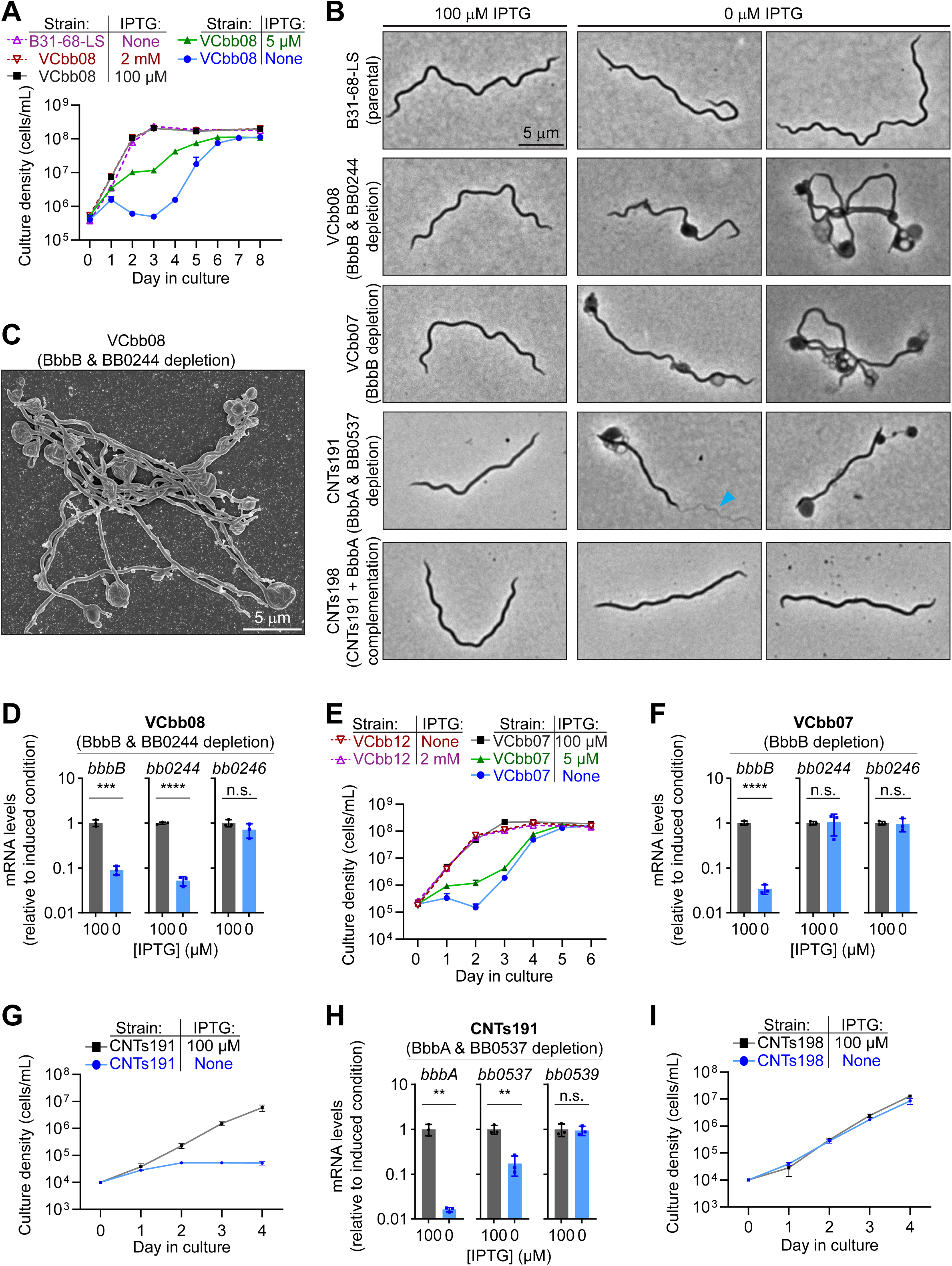
Characterization of growth, morphology, and gene expression in BbbB and BbbA depletion strains. **A.** Growth curves of parental strain B31-68-LS (violet) and derived strain VCbb08 (containing IPTG-inducible *bbbB* and *bb0244*) grown with the IPTG amounts indicated at the right. Shown are means ± standard deviations of results from *n*=3 cultures. **B.** Phase contrast light micrographs of cells of control and bactofilin depletion strains grown with 100 μM IPTG (left) or for 48 h without IPTG (0 μM IPTG, right). The strain identities are provided at the left. One image is provided for growth with IPTG, and two for growth without. The blue arrowhead points to flagella released from the periplasmic space in a cell of strain CNTs191 after 48 h without IPTG. **C.** Scanning electron micrograph of cells of strain VCbb08 grown for two days without IPTG. **D.** Gene expression (mRNA) levels for *bbbB*, *bb0244*, and *bb0246* measured in strain VCbb08 grown with 100 μM IPTG or in the absence of IPTG for 24 hours. Expression (individual values as well as means ± standard deviations from *n*=3 cultures) of each gene is shown relative to the mean value for the same gene in the culture grown in the presence of IPTG. **E.** Growth curves of strains VCbb12 (expressing its native *bbbB* gene and also carrying a shuttle vector-borne, IPTG-inducible *bbbB* copy) and VCbb07 (carrying only a shuttle vector-borne, IPTG-inducible *bbbB* copy) grown with the IPTG amounts indicated at the right. Shown are means ± standard deviations of results from *n*=3 cultures. **F.** Gene expression (mRNA) levels for *bbbB*, *bb0244*, and *bb0246* measured in strain VCbb07 grown with 100 μM IPTG or in the absence of IPTG for 24 hours. Expression (individual values as well as means ± standard deviations from *n*=3 cultures) of each gene is shown relative to the mean value for the same gene in the culture grown in the presence of IPTG. **G.** Growth curves of strain CNTs191 (containing IPTG-inducible *bbbA* and *bb0537*) grown with or without IPTG. Shown are means ± standard deviations of results from *n*=3 cultures. **H.** Gene expression (mRNA) levels for *bbbA*, *bb0537*, and *bb0539* measured in strain CNTs191 grown with 100 μM IPTG or in the absence of IPTG for 24 hours. Expression (individual values as well as means ± standard deviations from *n*=3 cultures) of each gene is shown relative to the mean value for the same gene in the culture grown in the presence of IPTG. **I.** Growth curves of strain CNTs198 (conditional *bbbA* and *bb0537* expression strain complemented with shuttle vector-borne, constitutively expressed *bbbA*) grown with or without IPTG. Shown are means ± standard deviations of results from *n*=3 cultures. **D**, **F**, and **H**: *p*-values (two-tailed, unpaired *t* test): ****, *p*<0.0001; ***, *p*<0.001; **, *p*<0.01; n.s., not significant, *p*>0.05.

Incubating VCbb08 without IPTG for 24 hours decreased transcript levels for *bbbB* and the downstream gene *bb0244* by more than 90%, while expression of the upstream gene *bb0246* (Fig. S1) was unaffected (Fig. 3D). This indicates that *bbbB* and *bb0244* form an operon and that depletion of either protein could be responsible for the growth and morphology defects observed (Figs. 3A-C). To distinguish between the functions of *bbbB* and *bb0244*, we first introduced a shuttle vector carrying an IPTG-inducible copy of *bbbB* into strain B31-68-LS (Fig. S3). The resulting strain, VCbb12, displayed similar growth kinetics in the presence or absence of 100 µM IPTG (Fig. 3E). In this background we deleted the chromosomal *bbbB* copy, yielding strain VCbb07 that carries its sole *bbbB* copy on the resident shuttle vector under IPTG-inducible control (Fig. S3). Strain VCbb07 displayed wild-type growth kinetics in the presence of 100 μM IPTG, while its growth was impaired at 5 or 0 μM IPTG, respectively (Fig. 3E). As with strain VCbb08, growth of strain VCbb07 was eventually observed after prolonged incubation without IPTG (Fig. 3E). Culturing strain VCbb07 in the absence of IPTG lowered *bbbB* transcript levels within 24 hours (Fig. 3F) and caused blebbing within 48 hours (Fig. 3B), but did not affect *bb0244* and *bb0246* transcript levels (Fig. 3F). Presumably, *bb0244* is transcribed in this strain via readthrough from the upstream *aacC1* gene (Fig. S3), which is controlled by the constitutive P*_flgB_* promoter (103). Taken together, our results indicate that BbbB is essential for *B. burgdorferi* growth in culture and maintenance of the cells’ normal morphology.

### BbbA is also required for *B. burgdorferi* growth in culture and maintenance of normal cell morphology

To investigate BbbA, we created depletion strain CNTs191 by placing the native *bbbA* (*bb0538*) gene under the control of the synthetic IPTG-inducible promoter P*_pQE30_* (101) (Fig. S3). Like strains VCbb08 and VCbb07, we derived CNTs191 from the LacI-expressing parent strain B31-68-LS (Fig. S3). CNTs191 exhibited a profound growth defect upon withdrawal of IPTG from the culture medium (Fig. 3G). By two days of *bbbA* depletion, the cells also accumulated severe morphological defects, including membrane blebbing (Fig. 3B) and release of periplasmic flagella (Fig. 3B, arrowhead). Within 24 hours of IPTG withdrawal, transcripts levels for *bbbA* and the downstream gene *bb0537* were reduced by 98% and 83%, respectively (Fig. 3H), suggesting that depletion of *bbbA* had polar effects. Meanwhile, expression of the upstream gene *bb0539* (Fig. S1) was unaffected by IPTG withdrawal (Fig. 3H). Additionally, the growth and cell morphology defects caused by *bbbA* (and *bb0537*) depletion were fully complemented (Figs. 3B,I) in strain CNTs198, which carries a second *bbbA* copy expressed constitutively from a shuttle vector (Fig. S3). These results indicate that, like *bbbB*, *bbbA* is also essential for cell replication and morphology maintenance in *B. burgdorferi*.

### Fluorescent protein-tagged BbbA and BbbB localize to known zones of *B. burgdorferi* cell growth

To further investigate BbbA and BbbB, we determined their subcellular localization by expressing mCherry-BbbA and mCherry-BbbB, respectively, from shuttle vectors (see strains CNTs216 and CJW_Bb176 in Fig. S3). The mCherry signal in both mCherry-BbbB and mCherry-BbbA-expressing cells concentrated at mid-cell (Figs. 4A,B). Demograph analyses of populations of cells confirmed mid-cell accumulation of the mCherry signal, which was clearest in spirochetes belonging to the middle ∼60% of the cell length distribution (Figs. 4C,D). In some of the longer cells we also observed mCherry-BbbB signal accumulation at the one-quarter and three-quarter positions along the cell length (Figs. 4C,E), in a pattern reminiscent of new peptidoglycan insertion sites (93). Western blotting of cell lysates obtained from these strains with an anti-mCherry antibody revealed both the full length mCherry-bactofilin fusions (mCherry-BbbA: expected ∼43 kDa, mCherry-BbbB: expected ∼48 kDa), as well as a smaller protein similar in size to free mCherry (∼27 kDa) (Fig. S4A). This result suggests that the mCherry-bactofilin constructs undergo processing within the bactofilin N-terminal tail, a phenomenon previously observed in *Myxococcus xanthus* and *Proteus mirabilis* (104, 105), or within the mCherry-bactofilin linker.

**Figure 4.**
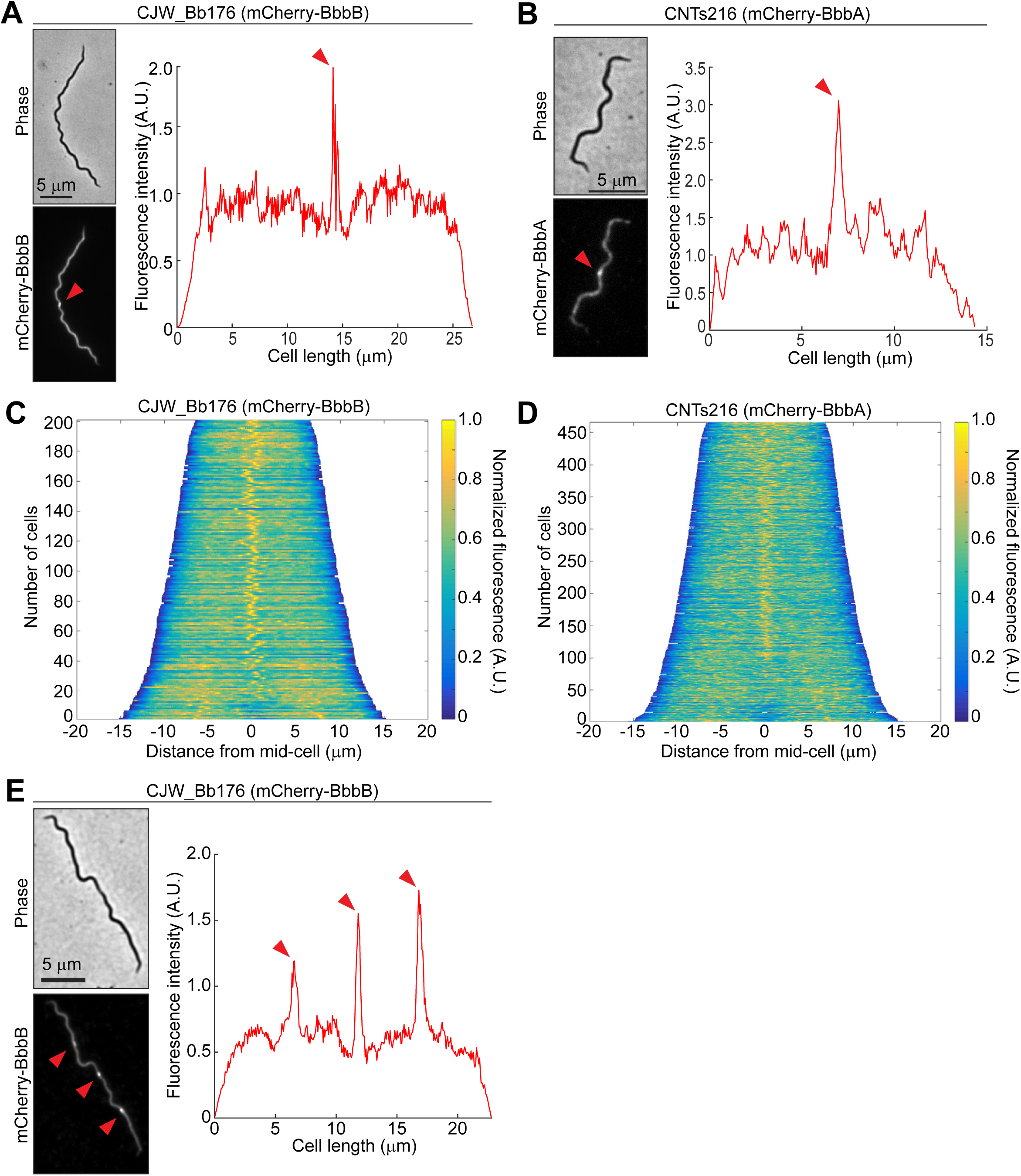
Fluorescence localization in strains expressing mCherry-tagged bactofilins. **A.** Phase contrast and fluorescence micrographs (left), and line intensity profiles (right) of a cell of strain CJW_Bb176 expressing mCherry-BbbB. Mid-cell signal accumulation is indicated by red arrowheads. A.U., arbitrary units. **B.** Same as in A but for a cell of strain CNTs216 expressing mCherry-BbbA. **C.** Demograph analysis of a population of 201 cells imaged from an exponentially growing culture of strain CJW_Bb176. Cells are arrayed from the shortest at the top to the longest at the bottom and are aligned along their mid-cell positions. The color of each line reflects the distribution of the mCherry signal along the cell length in a single cell, according to the color scale at the right. A.U., arbitrary units. **D.** Same as in C but for a population of 477 cells from strain CNTs216. A.U., arbitrary units. **E.** Same as in A but for a longer cell of CJW_Bb176 with additional fluorescence accumulation at quarter and three-quarter locations along the cell length, as indicated by red arrowheads.

To exclude the possibility that the mCherry-containing cleavage products accumulate at mid-cell, we imaged cells expressing free mCherry, mCherry fused to the N-terminal peptide of BbbB (mCherry-BbbB_1-21_), and mCherry fused to the first half of this peptide (mCherry-BbbB_1-10_). These constructs all displayed similar, mostly uniform mCherry signal throughout the cell length (Fig. S5), with the exception of reduced signal at the poles and the division site of the longest cells, which often contain periplasm and little to no cytoplasm (42). We also observed small dips in mCherry signal flanking mid-cell in pre-divisional cells (Fig. S5), likely representing the cytosolic constriction that occurs at sites of new flagellar basal body insertion (42). These observations indicate that the accumulation of fluorescence signal at mid-cell, as well as the one-quarter and three-quarter cell positions signal in mCherry-bactofilin-expressing strains CJW_Bb176 and CNTs216 cannot be attributed to mCherry-containing cleaved forms of the mCherry-tagged bactofilins.

### BbbB localization requires BbbA

Given the similar *bbbB* and *bbbA* depletion phenotypes (Figs. 3A-H) and same pattern of mid-cell localization observed for both mCherry-bactofilin fusion proteins (Fig. 4), we investigated whether bactofilins BbbA and BbbB function in the same pathway. To do so, we expressed mCherry-BbbB in the *bbbA* depletion background (strain CNTs220, Fig. S3). The resulting strain, CNTs220, exhibited mid-cell accumulation of mCherry signal when BbbA was expressed during growth in the presence of IPTG (Figs. 5A,B). In contrast, BbbA depletion led to delocalization of the mCherry-BbbB signal, which became dispersed throughout the cell body, except at the poles and future division site (Figs. 5C,D). Unlike with the free mCherry or mCherry-BbbB N-terminal peptide fusion constructs, we did not observe mCherry-BbbB signal dips flanking the mid-cell (Figs. 5D and S5). This difference in localization may arise from physiological differences between BbbA-expressing (Fig. S5) and BbbA-depleted (Figs. 5C,D) cells, or from protein behavior differences between mCherry-BbbB and mCherry or mCherry-BbbB N-terminal peptide fusions. BbbA depletion did not notably affect the relative abundance of the cleaved mCherry compared to full length mCherry-BbbB (Fig. S4B). We conclude that BbbA is required for the observed mCherry-BbbB localization to mid-cell and the one-quarter and three-quarter cell positions.

**Figure 5.**
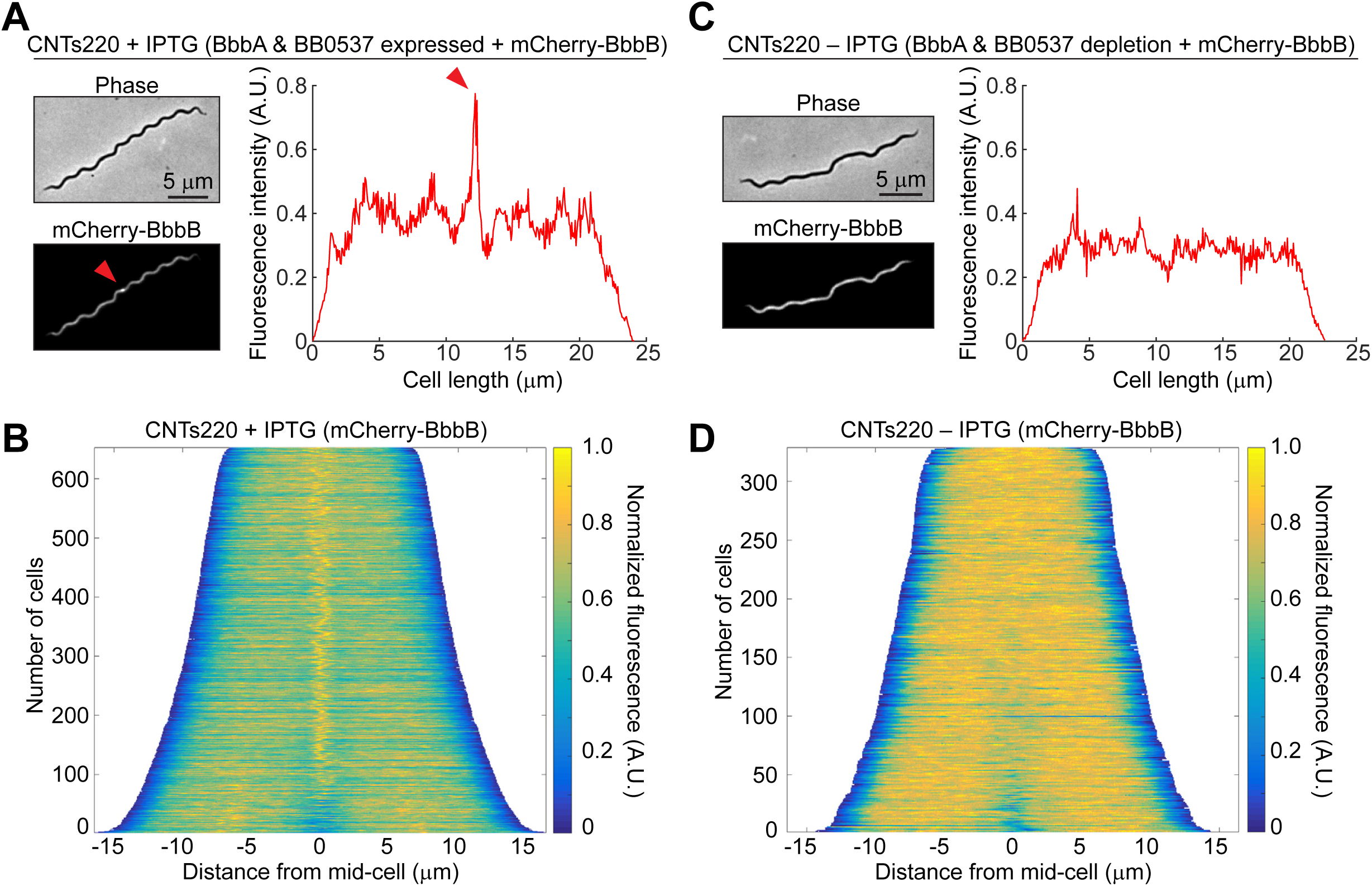
Role of BbbA in localizing mCherry-BbbB. **A.** Phase contrast and fluorescence micrographs (left), and line intensity profile (right) of a cell of strain CNTs220 (carrying IPTG-inducible *bbbA* and shuttle-vector borne constitutively expressed *mcherry-bbbB*) grown with 1 mM IPTG. Mid-cell accumulation of the mCherry signal is indicated by red arrowheads. A.U., arbitrary units. **B.** Demograph of a population of 653 cells of strain CNTs220 grown in the presence of IPTG. Cells are arrayed from the shortest at the top to the longest at the bottom and are aligned along their mid-cell positions. The color of each line reflects the distribution of the mCherry signal along the cell length, according to the color scale at the right. A.U., arbitrary units. **C.** Phase contrast and fluorescence micrographs (left), and line intensity profile (right) of a cell of strain CNTs220 grown in the absence of IPTG for 24 h. A.U., arbitrary units. **D.** Demograph of a population of 329 cells of strain CNTs220 grown in the absence of IPTG for 24 h. Cells are arrayed from the shortest at the top to the longest at the bottom and are aligned along their mid-cell positions. The color of each line reflects the distribution of the mCherry signal along the cell length, according to the color scale at the right. A.U., arbitrary units.

### BbbA and BbbB spatially organize peptidoglycan insertion in *B. burgdorferi*

The observation that mCherry-BbbB localizes (Figs. 4A,C,E) to zones of peptidoglycan insertion (93) in a BbbA-dependent manner (Fig. 5) prompted us to investigate the roles of BbbA and BbbB in peptidoglycan insertion. To this end, we pulse-labeled cells of the *bbbB* and *bbbA* depletion strains VCbb07 and CNTs191 (Fig. S3), respectively, with the fluorescent D-amino acid analog 7-hydroxycoumarin-amino-D-alanine (HADA), which labels new sites of peptidoglycan insertion in various bacteria, including *B. burgdorferi* (93, 106). When grown in the presence of IPTG to maintain bactofilin expression, both strains accumulated HADA signal predominantly at mid-cell (Figs. 6A-D). We also observed unipolar HADA signal accumulation in some, mostly shorter cells (Figs. 6B,D, arrowheads), consistent with incorporation of label during septal peptidoglycan synthesis and new pole formation when the preceding cell division occurred during the HADA labeling pulse, as previously shown (93). HADA accumulation at the one-quarter and three-quarter positions along the cell length could also be detected in some longer cells (Figs. 6E,F), consistent with the previous report (93). Using the same strains, bactofilin depletion for 24 hours prior to HADA labeling yielded cells with patchy fluorescence along most of the cell length, except at the poles (Figs. 6G-J). These patches of HADA signal were heterogeneous in size and intensity and were observed in both shorter and longer cells (Figs. 6G-J). Some mid-cell signal was still observed (Figs. 6G,I), likely due to incomplete depletion of the bactofilins over approximately three generations of IPTG-free growth. Since prolonged bactofilin depletion caused membrane blebbing (Figs. 3B,C), and the resulting cells were not amenable to washing or generation of the cell outlines required for quantitative microscopy image analysis, we were not able to reliably assess the effects of extended bactofilin depletion on peptidoglycan insertion. Nevertheless, our results implicate BbbB and BbbA in the localization of new peptidoglycan insertion in *B. burgdorferi*.

**Figure 6.**
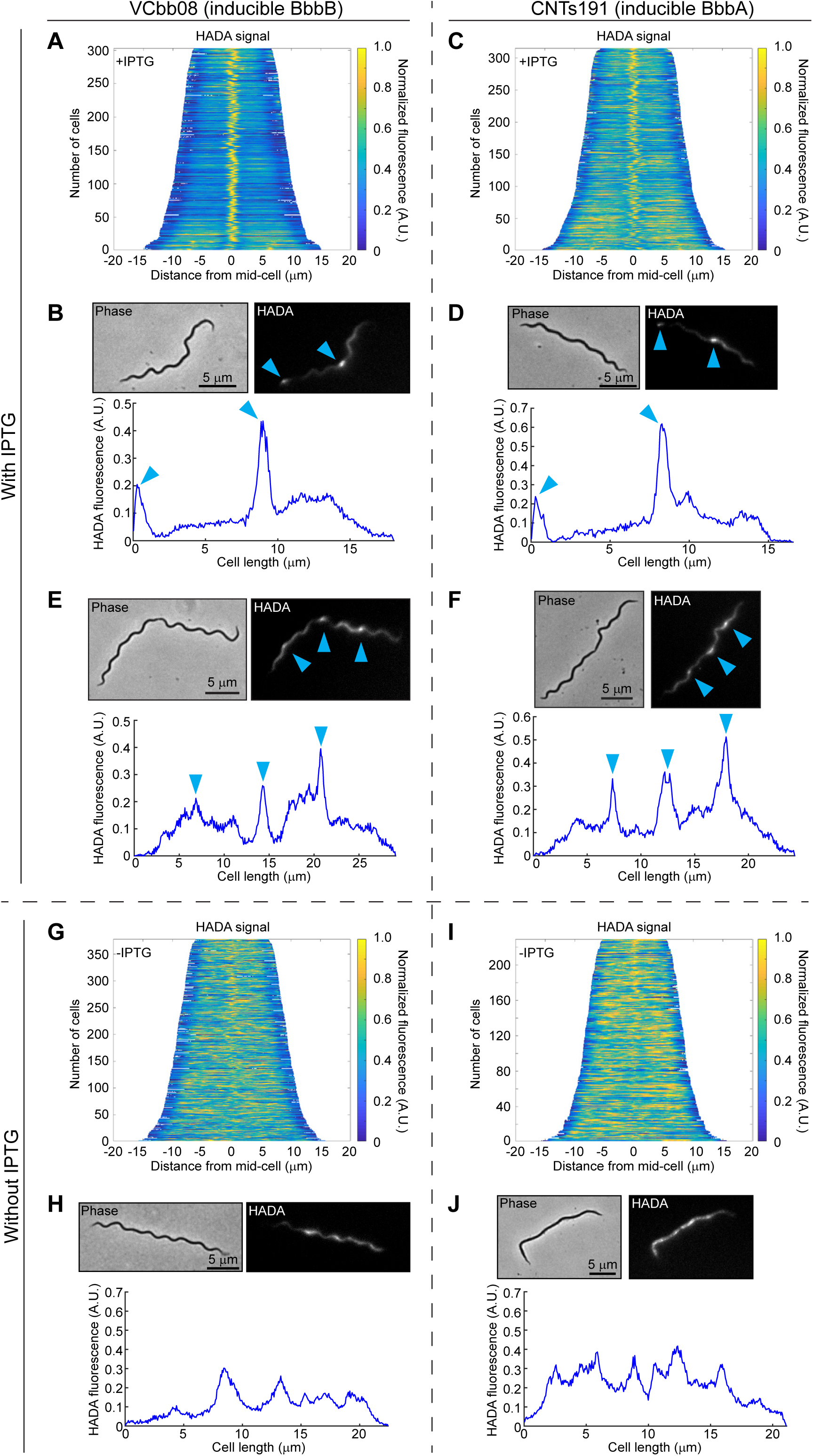
Effects of bactofilin depletion on the localization of new peptidoglycan insertion. **A.** Demograph of a population of 303 cells of strain VCbb08 (carrying IPTG-inducible *bbbB*) grown in the presence of IPTG and pulse-labeled with HADA for 60 min. Cells are arrayed from the shortest at the top to the longest at the bottom and are aligned along their mid-cell positions. The color of each line reflects the distribution of the HADA signal along the cell length in a single cell, according to the color scale at the right. A.U., arbitrary units. **B.** Phase contrast and fluorescence micrographs (top), and line intensity profile (bottom) of a cell from A. Mid-cell and polar accumulation are indicated by blue arrowheads. A.U., arbitrary units. **C.** Same as in A but for a population of 315 cells of strain CNTs191 (carrying IPTG-inducible *bbbA* and *bb0537*) grown in the presence of IPTG and pulse-labeled with HADA for 60 min. **D.** Phase contrast and fluorescence micrographs (top), and line intensity profile (bottom) of a cell from C. Mid-cell and polar accumulation are indicated by blue arrowheads. A.U., arbitrary units. **E.** Same as in B for an additional cell from A. Mid-cell, quarter, and three-quarter accumulation are indicated by blue arrowheads. **F.** Same as in D for an additional cell from C. Mid-cell, quarter, and three-quarter accumulation are indicated by blue arrowheads. **G.** Same as in A but for a population of 379 cells of strain VCbb08 grown in the absence of IPTG for 24 hours, then pulse-labeled with HADA for 60 min. **H.** Phase contrast and fluorescence micrographs (top), and line intensity profile (bottom) of a cell from G. A.U., arbitrary units. **I.** Same as in A but for a population of 229 cells of strain CNTs191 grown in the absence of IPTG for 24 hours, then pulse-labeled with HADA for 60 min. **J.** Phase contrast and fluorescence micrographs (top), and line intensity profile (bottom) of a cell from I. A.U., arbitrary units.

### Bactofilin accumulation precedes peptidoglycan insertion at zones of cell elongation

To further clarify the role of BbbB and BbbA in *B. burgdorferi* peptidoglycan insertion, we expressed each mCherry-tagged bactofilin in the corresponding bactofilin depletion strain, yielding strains CJW_Bb279 and CNTs217, respectively (Fig. S3). Population-wide localization analyses revealed that neither the pattern of tagged bactofilin localization nor that of HADA incorporation changed noticeably regardless of whether the native untagged bactofilins were expressed (+IPTG) or depleted (-IPTG) (Figs. 7A,B). Alongside our growth and morphology complementation results (Fig. S6), these observations strengthen our conclusion that BbbA and BbbB, and not the downstream-encoded proteins (BB0244 and BB0537, respectively), regulate peptidoglycan insertion and thus maintain cell integrity. Visualization of individual cells revealed colocalization of the tagged bactofilins with HADA accumulation at mid-cell (Figs. 7C-E). However, we also observed striking differences between the localization of mCherry-BbbB and the HADA signal: we found that mCherry-BbbB could localize to the one-quarter and three-quarter cell positions in the absence of noticeable HADA accumulation during the preceding hour of pulse (Fig. 7D). The simplest explanation for this result is that BbbB accumulates at zones of cell growth before it either recruits or confines the peptidoglycan synthesis machinery to these locations.

**Figure 7.**
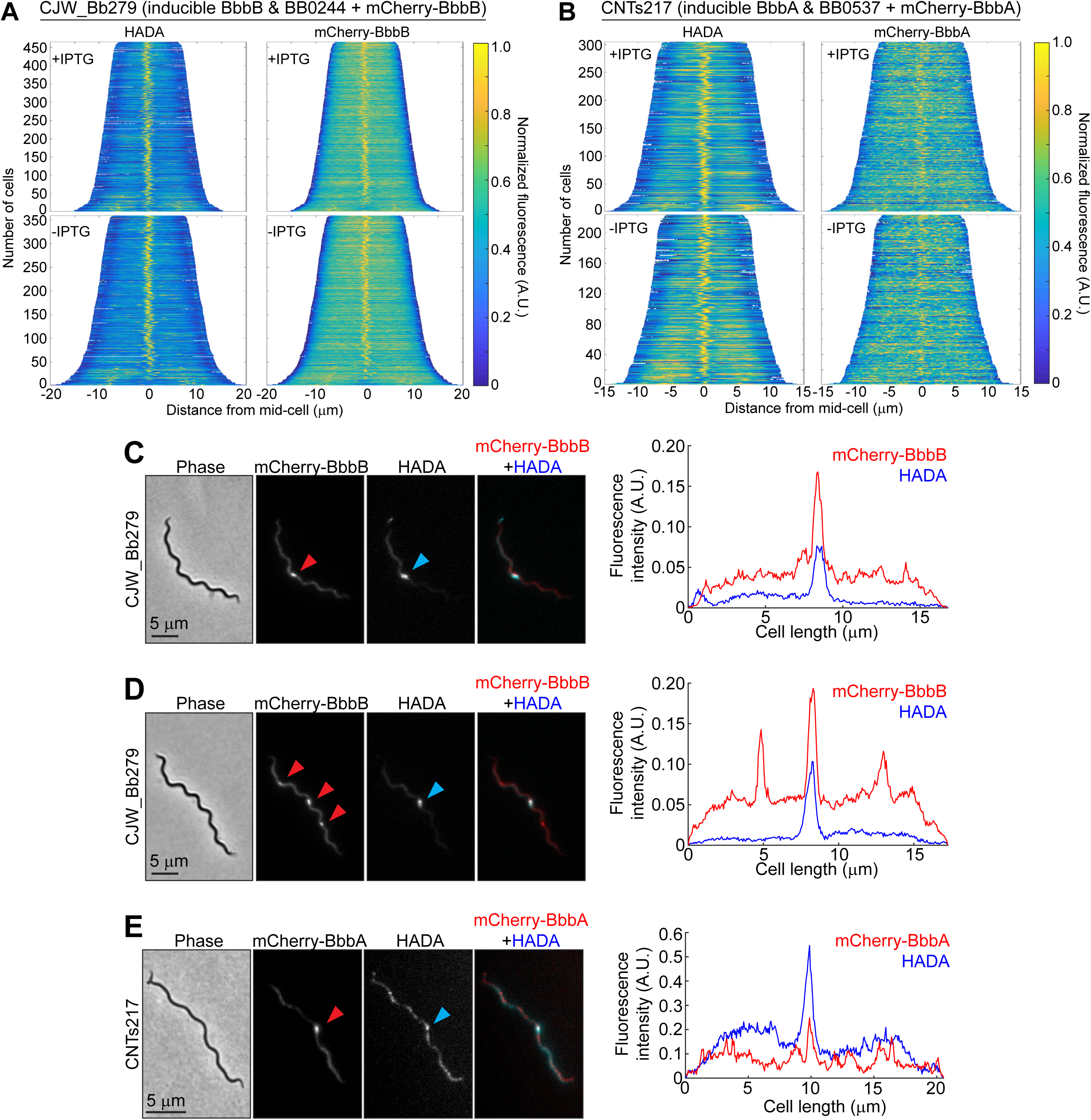
Role of BbbB and BbbA in the localization of new peptidoglycan insertion. **A.** Demographs of populations of cells of strain CJW_Bb279 (carrying IPTG-inducible *bbbB* and *bb0244* along with shuttle vector-borne constitutively expressed *mcherry-bbbB*) grown with 100 μM IPTG (top, *n* = 465 cells) or grown in the absence of IPTG for 24 h (bottom, *n* = 359 cells). The cells were labeled with HADA for 60 min. Demographs of the HADA signal are on the left, while those of the mCherry-BbbB signal are on the right. The cells are arrayed from the shortest at the top to the longest at the bottom and are aligned along their mid-cell positions. The color of each line reflects the distribution of the fluorescent signal along the cell length, according to the color scale at the right. A.U., arbitrary units. **B.** Same as in A but for strain CNTs217 (carrying IPTG-inducible *bbbA* and *bb0537* along with shuttle vector-borne constitutively expressed *mcherry-bbbA*) grown with 100 μM IPTG (top, *n* = 305 cells) or grown in the absence of IPTG for 24 h (bottom, *n* = 224 cells). The cells were labeled with HADA for 60 min. Demographs of the HADA signal are on the left, while those of the mCherry-BbbA signal are on the right. **C.** Phase contrast and fluorescence micrographs (left), and line intensity profiles (right) of a cell of strain CJW_Bb279 grown with 100 μM IPTG and labeled with HADA. From left to right, the images are: phase contrast micrograph; fluorescence micrograph of mCherry-BbbB; fluorescence micrograph of HADA; and overlayed image of the mCherry-BbbB signal (in red) with the HADA signal (in blue). Mid-cell accumulation of mCherry-BbbB and HADA signal are indicated by red and blue arrowheads, respectively. Right, fluorescence intensity profile along the cell length. The mCherry-BbbB signal is in red, while the HADA signal is in blue. A.U., arbitrary units. **D.** Same as in C for an additional cell of strain CJW_Bb279 grown with 100 µM IPTG. Mid-cell, quarter, and three-quarter accumulation of mCherry-BbbB and mid-cell accumulation of HADA signal are indicated by red and blue arrowheads, respectively. **E.** Same as in C for a cell of strain CNTs217 grown with 100 µM IPTG. Mid-cell accumulation of mCherry-BbbA and HADA signal are indicated by red and blue arrowheads, respectively.

## DISCUSSION

Here, we report an initial characterization of the function of *B. burgdorferi* bactofilins BbbA and BbbB. Our preliminary investigation of BbbC suggests that this bactofilin carries functions distinct from those of BbbA and BbbB. We will report the results of our BbbC studies in a separate publication. Our findings implicate BbbA and BbbB in maintenance of *B. burgdorferi* cell integrity. We show that BbbA and BbbB are essential for *B. burgdorferi* viability, as we were unable to delete or even silence the chromosomal copies of *B. burgdorferi* bactofilin genes *bbbA* and *bbbB* without expressing a second copy of the targeted gene in trans. In contrast, most bactofilins studied to date are not essential for bacterial growth and viability, as shown by the successful generation of the corresponding bactofilin gene deletion mutants (72, 76, 79, 83, 84, 86–88, 92, 105). In *Chlamydia trachomatis,* CRISPR interference was used to deplete bactofilin BacA_CT_ and reveal its role in cell size maintenance (82). In *Bacillus subtilis* strain PY79, the bactofilin-encoding genes *bacE* (*yhbE*) and *bacF* (*yhbF*) could not be deleted individually but could be deleted together (107), and single gene deletions could be transferred into this background from strain 168 (78).

We demonstrate that mCherry-BbbB requires BbbA for its accumulation at the one-quarter, mid-cell, and three-quarter positions (Fig. 5). mCherry-BbbA itself accumulates at mid-cell (Figs. 4B,D and 7B,E) but we have not observed clear mCherry-BbbA accumulation at the one-quarter and three-quarter positions. However, mCherry-BbbA is expressed at lower levels than mCherry-BbbB (Fig. S4A), which may render detection of the small amounts of mCherry-BbbA that we presume initiate bactofilin accumulation at one-quarter and three-quarter positions hard to distinguish from the mCherry-BbbA signal distributed throughout the rest of the cell (Figs. 4B,D and 7B,E). BbbA could directly recruit BbbB to these locations through co-polymerization of the two bactofilins into the same BbbA-initiated protofilament, or interactions between separate homo-polymeric filaments. Such modes of co-assembly have been previously invoked for other bactofilins (72, 79, 83). However, mCherry-BbbA does eventually accumulate at mid-cell to levels clearly detectable above the rest of the cellular signal, which suggests that BbbA assembles slower than BbbB. Our observations are also consistent with a model in which small amounts of BbbA initially arrive at the one-quarter and three-quarter positions, recruit additional factors that transform these locations into suitable hubs for faster BbbB accumulation, while slower BbbA accumulation proceeds in parallel.

Irrespective of the mechanistic details of BbbA and BbbB recruitment, both tagged bactofilins accumulate at mid-cell (Figs. 4A-D), where peptidoglycan insertion primarily occurs in *B. burgdorferi* (Figs. 6A-D) (93). Depletion of either BbbA or BbbB alters the pattern of peptidoglycan insertion from concentrated accumulation at mid-cell and quarter-cell positions to dispersed insertion along the cell length (Figs. 6G-J). These findings suggest that proper localization of peptidoglycan insertion in *B. burgdorferi* is achieved through the joint activities of both bactofilins. Joint regulation of a cellular process by several bactofilins has been previously described in *C. crescentus* (72), *M. xanthus* (79), and *Rhodomicrobium vannielii* (83).

The delay in new peptidoglycan insertion relative to the arrival of BbbB at zones of cell wall growth (Fig. 7D), along with our finding that BbbB’s recruitment to these zones requires BbbA (Fig. 5), suggests that these two bactofilins mark locations of future cell wall growth. The mechanism that initially recruits BbbA to these locations remains unknown. It is possible that a dedicated recruitment factor localizes there, and then recruits BbbA and BbbB, similar to how SpmX recruits bactofilin BacA to the site of future stalk synthesis in *Asticaccaulis biprosthecum* (76). In *H. pylori*, bactofilin CcmA is thought to recognize positive Gaussian curvature and direct peptidoglycan insertion along the long helical axis of this bacterium, establishing and maintaining helicity (92). A different mechanism may be at work in *L. biflexa*, where the cell’s helical pitch depends on bactofilin LbbD, as demonstrated by opposing pitch changes caused by *lbbD* overexpression and deletion, respectively (88). In *B. burgdorferi*, the pattern of peptidoglycan insertion is similar whether the cells retain their wild-type flat wave morphology or grow as straight rods after genetic ablation of the flagella through deletion of the flagellin gene *flaB* (50, 93). This implies that BbbA and BbbB filaments do not rely on membrane curvature to position themselves at specified locations along the cell length.

How might BbbA and BbbB direct peptidoglycan insertion to the primary zones of *B. burgdorferi* cell wall growth? A plausible scenario is direct recruitment of peptidoglycan synthases by BbbA and/or BbbB. Such a mechanism has precedent in *C. crescentus*, where bactofilin BacA recruits a bifunctional peptidoglycan synthase with transglycosylase and transpeptidase domains, PbpC, to the base of the growing stalk (72). If a similar mechanism were at work in *B. burgdorferi*, either recruitment or activation of the peptidoglycan synthase(s) must be delayed relative to bactofilin accumulation to account for the delay we observe between bactofilin arrival and new peptidoglycan insertion at the aforementioned zones (Fig. 7D). It is possible that this bactofilin-driven peptidoglycan synthesis activity is provided by the elongasome itself, which the bactofilins could recruit to their sites of subcellular accumulation. This idea is supported by the localized bulging observed at these sites during MreB or RodA depletion in *B. burgdorferi* (95), which implies that the elongasome functions at the same subcellular locales as the bactofilins. It also explains the delay between bactofilin accumulation and new peptidoglycan insertion, as elongasome assembly and activation for peptidoglycan synthesis is an ordered, multistep process (37). Control of the *B. burgdorferi* MreB-driven elongasome by BbbA and BbbB would represent an evolutionarily-divergent mechanism from those imposing spatial separation between bactofilin and elongasome activities in *H. pylori* (92) or between bactofilin activity and the MreB-driven, specialized divisome of *C. trachomatis* (82, 108–111).

In addition to spatial control of peptidoglycan insertion, BbbA and BbbB may also play a role in peptidoglycan degradation. A recent study revealed an evolutionarily-conserved pairing between bactofilins and M23 endopeptidases homologous to *Hyphomonas neptunium* LmdC (84). While not universal (82), this pairing was observed in several other bacteria, including several Borreliaceae species (76, 84, 86, 87, 91). The conserved pairing of homologous genes across species can suggest a functional relationship between the encoded proteins (112). In *B. burgdorferi*, the M23 endopeptidase is encoded by gene *bb0246*, which is located immediately upstream of *bbbB* (Fig. S1A). It is possible that BbbB-associated BB0246 locally degrades peptidoglycan, which could facilitate new peptidoglycan insertion.

Why are bactofilins BbbA and BbbB essential for cell growth and integrity in *B. burgdorferi*, while their counterparts in other bacteria appear largely dispensable? Peptidoglycan insertion continued in *B. burgdorferi* cells depleted of bactofilin for 24 hours without any obvious morphological defects (Figs. 6 and S7), while 24-hour depletion of essential elongasome components MreB and RodA resulted in cell bulging (95). By 48 hours, depletion of the bactofilins caused cell blebbing (Figs. 3B,C), which is clearly distinct from the cell bulging caused by MreB or RodA depletion (95). These divergent effects suggest that the elongasome is still active in bactofilin-depleted *B. burgdorferi* cells. The cell envelope instability secondary to bactofilin depletion may therefore be due to uncoupling of elongasome-directed peptidoglycan synthesis from BB0246-mediated peptidoglycan degradation. This may generate areas of weakened cell wall, where the envelope becomes unstable and blebs. In contrast, control of BB0246-mediated peptidoglycan degradation and elongasome-directed peptidoglycan insertion by the same cytoskeletal bactofilin structure would ensure that the two activities occurred in the same location, thus preserving a regular pattern of growth, cellular integrity, and, by extension, ensuring viability.

## MATERIALS AND METHODS

### Strains and growth conditions

The *B. burgdorferi* strains used in this study are listed in Table 1. They were grown in complete Barbour-Stoenner-Kelly (BSK)-II medium at 34°C and under 5% CO_2_ atmosphere (113–115). The BSK-II medium contained 50 g/L bovine serum albumin (Millipore cat. 810036), 9.7 g/L CMRL-1066 (US Biological cat. 5900-01), 5 g/L neopeptone (Difco cat. 211681), 2 g/L yeastolate (Difco cat. 255772), 6 g/L HEPES (Millipore cat. 391338 or Fisher cat. BP310), 5 g/L Glucose (Sigma cat. G7021 or Fisher cat. BP350), 0.7 g/L sodium citrate (Sigma cat. C7254 or Fisher cat. BP327), 0.8 g/L sodium pyruvate (Sigma cat. P5280 or Fisher cat. BP356), 0.4 g/L N-acetylglucosamine (Sigma cat. A3286 or Thermo Scientific cat. A13047.09), 2.2 g/L sodium bicarbonate (Sigma cat. S3817 or S5761 or Fisher cat. BP328), and 60 mL/L heat-inactivated rabbit serum (Gibco cat. 161120). The pH of the medium was adjusted to 7.60 with 5 M sodium hydroxide. Occasionally, the medium was supplemented with 10 g/L gelatin. To avoid loss of endogenous plasmids in stationary phase (29), cultures were maintained in exponential growth, with passages performed at densities below 5x10^7^ cells/mL, unless otherwise indicated. Cell density was measured by direct counting of samples placed in disposable hemocytometers and observed on a microscope using darkfield illumination, as described before (116). As needed, isopropyl-β-D-1-thiogalactopyranoside (IPTG, Fisher cat. BP1755 or American Bioanalytical cat. AB00841) was added from a 1M stock in water to make the indicated final concentrations. Final antibiotic concentrations were: streptomycin, 100 μg/mL (117), gentamicin, 40 μg/mL (118), kanamycin, 200 μg/mL (119), and blasticidin S, 10 μg/mL (116).

**Table 1.** *B. burgdorferi* strains used in this study.

| Strain name | Record number | Relevant genotype | Growth conditions<sup>a</sup> | Source or reference |
| --- | --- | --- | --- | --- |
| S9 | CNTs005 | B31-A3-68 $\Delta bbe02::P_{flaB}$-*aadA* | Sm | (141) |
| B31-68-LS | CNTs017 | B31-A3-69 $\Delta bbe02::[P_{flgB}$-*lacI* $P_{flgB}$-*aadA*] | Sm | (102) |
| VCbb08 | CNTs173 | B31-68-LS *bbbB*::$[P_{flaB}$-*aacC1* $P_{flac}$-*bbbB*] | Sm, IPTG | This study |
| VCbb12 | CNTs177 | B31-68-LS / pBSV2_ $P_{flac}$-*BbbB* | Km, Sm | This study |
| VCbb07 | CNTs172 | B31-68-LS $\Delta bbbB::P_{flgB}$-*aacC1* / pBSV2_ $P_{flac}$-*BbbB* | Km, Sm, IPTG | This study |
| CJW_Bb176 | CNTs155 | S9 / pBSV2_ $P_{0826}$-*mCherry*<sup>Bb</sup>-*BbbB* | Km, Sm | This study |
| CJW_Bb185 | CNTs152 | S9 / pBSV2_ $P_{0826}$-*mCherry*<sup>Bb</sup>-*BbbA* | Km, Sm | This study |
| CJW_Bb279 | CNTs156 | VCbb08 / pBSV2_ $P_{0826}$-*mCherry*<sup>Bb</sup>-*BbbB*_ $P_{0026}$-*RBS-mEGFP*<sup>Bb</sup> | Km, Sm, IPTG | This study |
| CNTs191 | CNTs191 | B31-68-LS $\Delta bbbA::[r(P_{flaB}$-*aphI*<sup>Bb</sup>)_ $P_{pQE30}$-*bbbA*] | Km, Sm, IPTG | This study |
| CNTs198 | CNTs198 | CNTs191 / pBSV2G_ $P_{0526}$-*BbbA* | Sm, Gm, IPTG | This study |
| CNTs216 | CNTs216 | S9 / pBSV2G_ $P_{0526}$-*mCherry*<sup>Bb</sup>-*BbbA* | Sm, Gm | This study |
| CNTs217 | CNTs217 | CNTs191 / pBSV2G_ $P_{0526}$-*mCherry*<sup>Bb</sup>-*BbbA* | Sm, Gm, IPTG | This study |
| CNTs220 | CNTs220 | CNTs191 / pBSV2B_ $P_{0826}$-*mCherry*<sup>Bb</sup>-*BbbB* | Sm, Bs, IPTG | This study |
| CJW_Bb181 | CNTs148 | S9 / pBSV2G_ $P_{0826}$-*mCherry*<sup>Bb</sup> | Sm, Gm | This study |
| CNTs287 | CNTs287 | S9 / pBSV2B_ $P_{0826}$-*mCherry*<sup>Bb</sup>-*BbbB*<sub>1-10</sub> | Sm, Bs | This study |
| CNTs288 | CNTs288 | S9 / pBSV2B_ $P_{0826}$-*mCherry*<sup>Bb</sup>-*BbbB*<sub>1-21</sub> | Sm, Bs | This study |

<sup>a</sup>Sm, streptomycin; Km, kanamycin; Gm, gentamicin; Bs, blasticidin S.

The *E. coli* strains used in this study were propagated at 30°C on Luria Bertani agar plates or in Super Broth (35 g/L bacto-tryptone, 20 g/L yeast extract, 5 g/L sodium chloride, 6 mM sodium hydroxide) liquid cultures, with shaking. Antibiotics were used as follows: gentamicin at 15 to 20 μg/mL, streptomycin or spectinomycin at 50 μg/mL, rifampicin at 50 μg/mL (plate) or 25 μg/mL (liquid culture), and kanamycin at 50 μg/mL. Most often, NEB 5-alpha (New England Biolabs, cat. C2987) was used as a host for genetic manipulations and archiving constructs, except that plasmids containing IPTG-inducible promoters were generated and maintained in NEB 5-alpha F’ *Iq* hosts (New England Biolabs, cat. C2992) (95). Transformants were obtained by heat shock, then allowed to recover for one hour with shaking at 30°C in SOC medium (20 g/L tryptone, 5 g/L yeast extract, 10 mM NaCl, 2.5 mM KCl, 10 mM MgSO_4_, and 20 mM glucose) or Super Broth before plating.

### Plasmid construction methods

The plasmids used in this study are listed in Table 2 and the sequences of the oligonucleotide primers used in their construction are provided in Table 3. The construction methods are described below. Standard molecular biology techniques were used unless otherwise indicated. TOPO Cloning kits were purchased from Invitrogen. Restriction endonucleases were purchased from New England Biolabs.

**Table 2.**
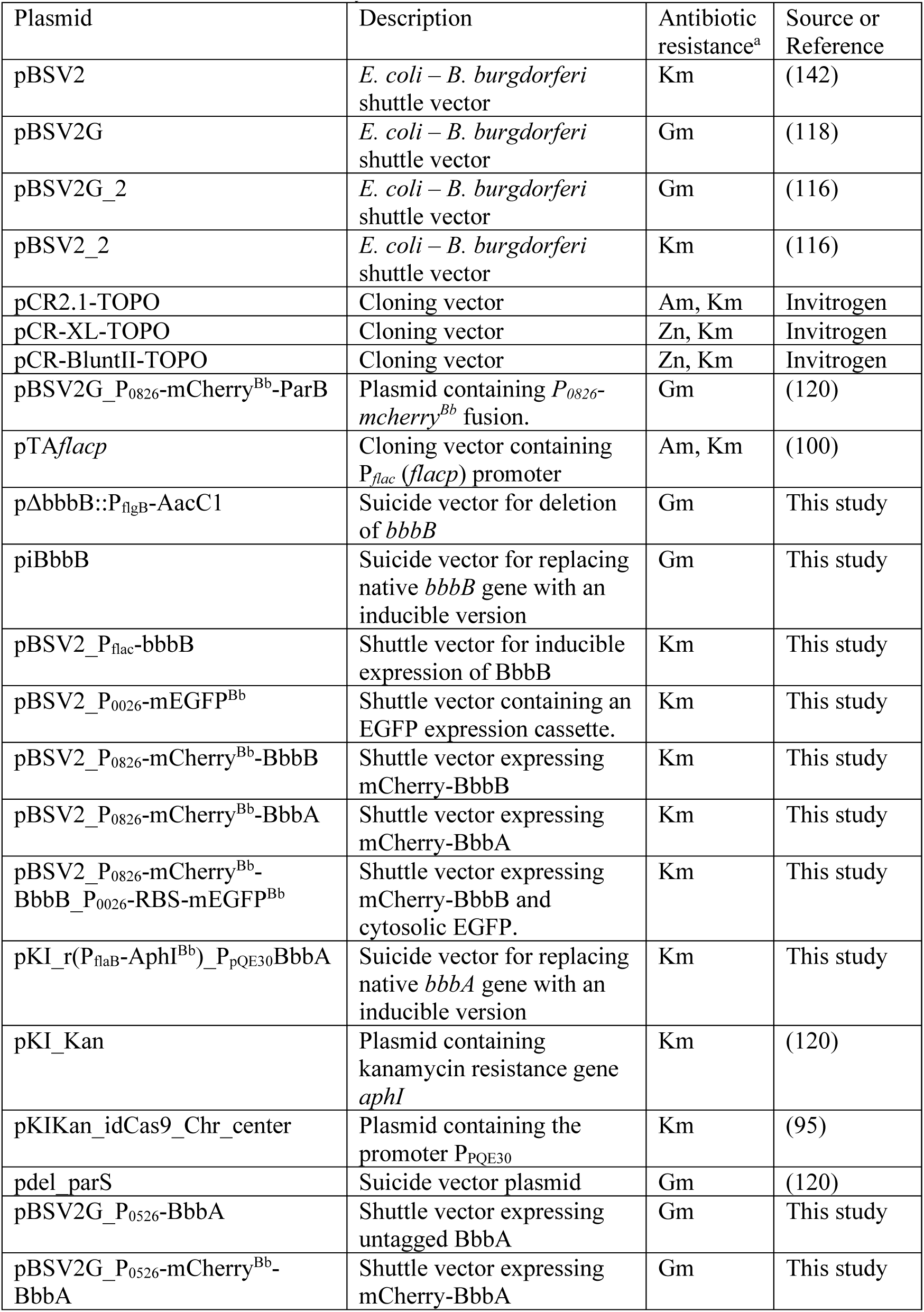
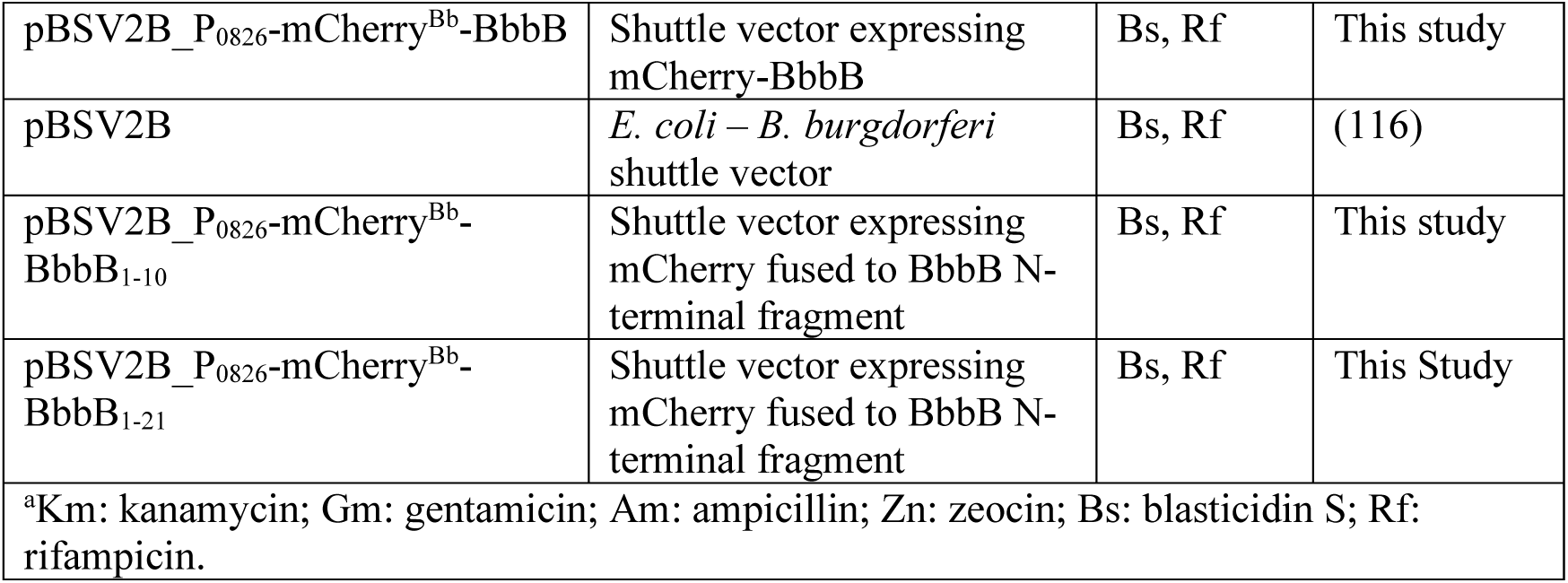
Plasmids used in this study.

**Table 3.**
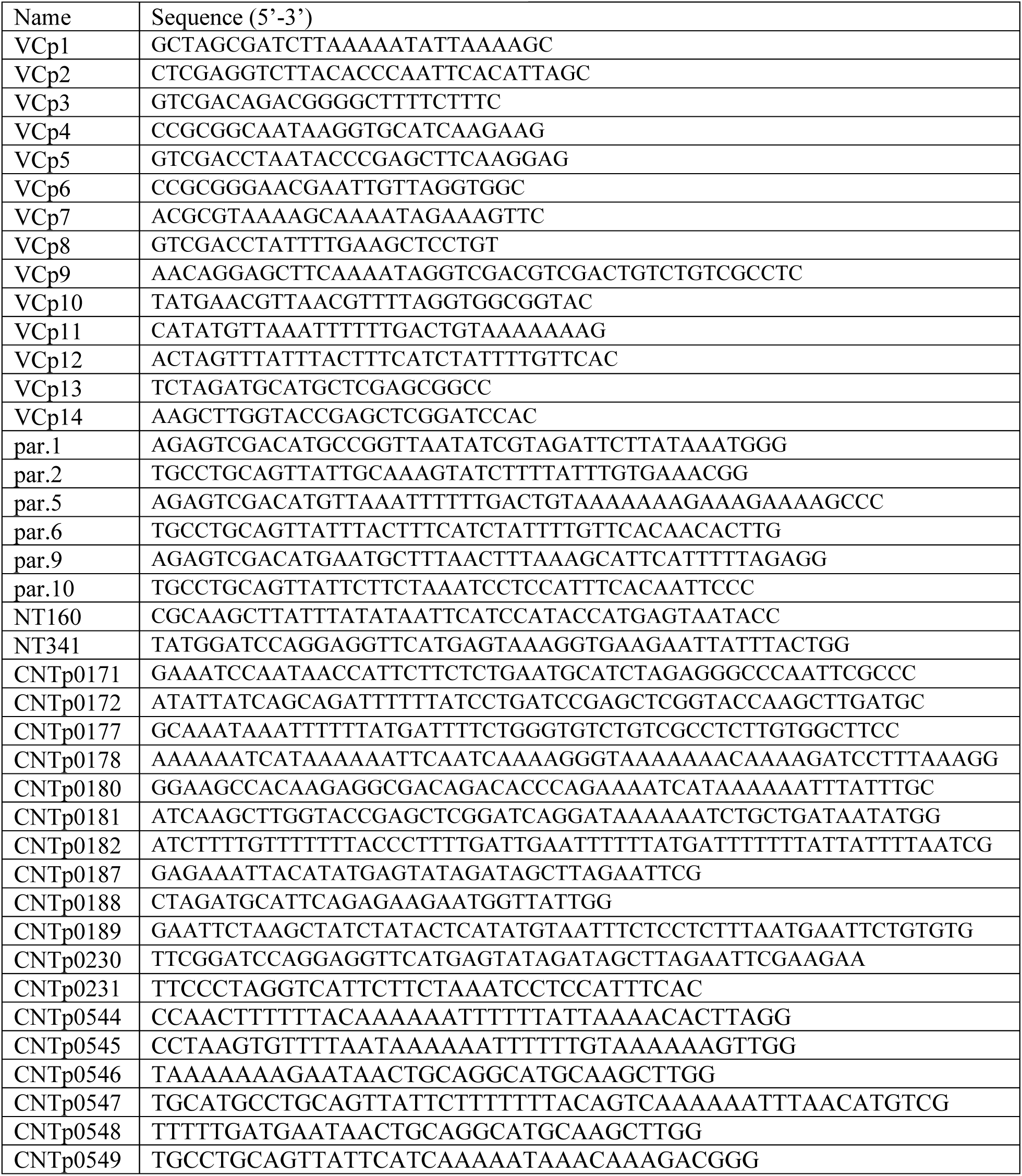
Oligonucleotide primers used in this study.

#### pΔbbbB::P_flgB_-AacC1

A fragment of the *B. burgdorferi* chromosome was PCR-amplified with primers VCp1 and VCp2 and cloned into plasmid pCR2.1-TOPO. In the resulting intermediate plasmid, inverse PCR was performed with primers VCp3 and VCp4 to delete the central portion of gene *bbbB* (*bb0245*). The product was subcloned into pCR-XL-TOPO. The resulting plasmid was then digested with SalI and SacII to release the pCR2.1-TOPOΔbbbB backbone. The P*_flgB_-aacC1* gentamicin resistance cassette was PCR amplified from pBSV2G using VCp5 and VCp6, subcloned into pCR-BluntII-TOPO, then released as a SalI/SacII fragment and ligated with the SalI/SacII pCR2.1-TOPOΔbbbB backbone described above. The product was digested with AclI and then re-ligated to inactivate the ampicillin resistance gene, yielding plasmid pΔbbbB::P_flgB_-AacC1.

#### piBbbB

A fragment of gene *bb0246* was PCR-amplified using primers VCp7 and VCp8 and cloned into pCR-BluntII-TOPO. A P*_flaB_-aacC1* gentamicin resistance cassette previously described (102) was PCR-amplified using primers VCp9 and VCp10 and cloned into pCR-BluntII-TOPO. The two plasmids were digested with SalI and SpeI and the P*_flaB_-aacC1* fragment was ligated to the pCR-BluntII-TOPO_bb0246 backbone. The *bbbB* sequence was PCR-amplified using primers VCp11 and VCp12 and cloned into pCR-BluntII-TOPO, then *bbbB* was released as a NdeI/SpeI fragment and cloned into the NdeI/SpeI-digested pTA*flacp* (100). The resulting P*_flac_*-*bbbB* fragment was released as an XbaI/KpnI fragment and ligated to the SpeI/KpnI backbone of pCR-BluntII-TOPO_bb0246_P_flaB_-aacC1 to yield plasmid piBbbB.

#### pBSV2_P_flac_-BbbB

A P*_flac_*-*bbbB* fragment described above was PCR-amplified using VCp13 and VCp14 and cloned in pCR-BluntII-TOPO, then released as a BamHI/XbaI fragment, and finally cloned into BamHI/XbaI-digested pBSV2. pBSV2_P_0826_-mCherry^Bb^-BbbB and pBSV2_P_0826_-mCherry^Bb^-BbbA. Through intermediate constructs, the following fragments were assembled in the pBSV2_2 backbone. A P*_0826_*-*mcherry^Bb^* transcriptional fusion lacking a STOP codon was previously described in plasmid pBSV2G_P_0826_-mCherry^Bb^-ParB (120) and was inserted between the SacI and KpnI sites of pBSV2_2. The bactofilin genes were amplified using primers par.5 and par.6 (*bb0245*/*bbbB*), or par.9 and par.10 (*bb0538*/*bbbA*). The resulting fragments were digested with SalI and PstI and inserted into the corresponding sites of pBSV2_2. The mCherry-bactofilin fusions thus generated contain a GYRSSRVD linker, with the mCherry-BbbA linker including the 14 amino acids preceding the likely BbbA translational start site (Fig. S2B).

#### pBSV2_P_0826_-mCherry^Bb^-BbbB_P_0026_-RBS-mEGFP^Bb^

The following steps were taken to generate pBSV2_P_0026_-RBS-mEGFP^Bb^: promoter P*_0026_* (116), flanked by SacI and BamHI sites, and *megfp^Bb^* (116), PCR-amplified using primers NT160 and NT341 and digested with BamHI and HindIII, were assembled between the SacI and HindIII sites of plasmid pBSV2_2 (116). A fragment containing the P*_0026_*-*RBS-megfp^Bb^* sequence was released from pBSV2_P_0026_-RBS-mEGFP^Bb^ as a BsrBI/AvrII fragment and then ligated into the FspI/AvrII backbone of pBSV2_P_0826_-mCherry^Bb^-BbbB.

#### pKI_r(P_flaB_-aphI)_P_pQE30_-BbbA

A region of the *B. burgdorferi* chromosome upstream of *bbbA*, encompassing nucleotides 550,492 to 548,886, was PCR-amplified with primers CNTp0181 and CNTp0182. A region of the *B. burgdorferi* chromosome, encompassing nucleotides 548,843 to 547,208 and containing *bbbA* (beginning at the second annotated start site, Fig. S2B), and downstream genes were PCR-amplified with primers CNTp0187 and CNTp0188. Both amplification reactions were done using genomic DNA from strain CJW_Bb378 (120). The antibiotic resistance cassette P*_flaB_-aphI* was amplified from pKI_Kan (120) using primers CNTp0178 and CNTp0177. The synthetic promoter P*_pQE30_* was amplified from plasmid pKIKan_idCas9_Chr_center (95) using primers CNTp0180 and CNTp0189. The backbone was amplified using primers CNTp0171 and CNTp0172 from a derivative of pdel_parS (120). Fragments were assembled using the NEBuilder HiFi assembly kit.

#### pBSV2G_P_0526_-BbbA

The gene *bbbA* (beginning at the second annotated start site, Fig. S2B) was amplified from B31-68-LS using primers CNTp0230 and CNTp0231 and the resulting PCR product was digested with BamHI and AvrII. Promoter P*_0526_* was previously described (Takacs et al 2018) and is flanked by SacI and BamHI sites. Through intermediary constructs, the SacI/BamHI P*_0526_* fragment and the BamHI/AvrII *bbbA* fragment were assembled between the SacI/AvrII sites of pBSV2G_2 (116).

#### pBSV2G_P_0526_-mCherry^Bb^-BbbA

*mcherry^Bb^-BbbA* was released from pBSV2_P_0826_-mCherry^Bb^-BbbA with BamHI-HF and AvrII and ligated into the BamHI/AvrII-linearized backbone of pBSV2G_P_0526_-BbbA.

#### pBSV2B_P_0826_-mCherry^Bb^-BbbB

P*_0826_-mcherry^Bb^-BbbB* was released from plasmid pBSV2_P_0826_-mCherry^Bb^-BbbB with BsrBI and AvrII and ligated into the BsrBI/AvrII-linearized backbone of pBSV2B (116).

#### pBSV2B_P_0826_-mCherry^Bb^-BbbB_1-10_

A fragment of pBSV2B_P_0826_-mCherry^Bb^-BbbB including the first 30 nucleotides of *bbbB* was PCR amplified with primers CNTp0544 and CNTp0547. A second fragment of pBSV2B_P_0826_-mCherry^Bb^-BbbB containing the stop codon of *bbbB* was PCR amplified using primers CNTp0545 and CNTp0546. These fragments were assembled using the NEBuilder HiFi assembly kit.

#### pBSV2B_P_0826_-mCherry^Bb^-BbbB_1-21_

A fragment of pBSV2B_P_0826_-mCherry^Bb^-BbbB including the first 63 nucleotides of *bbbB* was PCR-amplified with primers CNTp0544 and CNTp0549. A second fragment of pBSV2B_P_0826_-mCherry^Bb^-BbbB containing the stop codon of *bbbB* was PCR amplified using primers CNTp0545 and CNTp0548. These fragments were assembled using the NEBuilder HiFi assembly kit.

### *B. burgdorferi* strain generation

*B. burgdorferi* competent cells were prepared as described before (121). Cultures were allowed to grow to densities between 2x10^7^ cells/mL and 10^8^ cells/mL. The cells were centrifuged for 10 min at 10,000 x g and 4°C and resuspended in ice-cold electroporation solution (EPS), which contained 93.1 g/L sucrose (American Bioanalytical cat. AB01900) and 150 mL/L glycerol (American Bioanalytical cat. AB00751). A total of three centrifugation steps followed by resuspension in decreasing amounts of EPS achieved the desired salt removal and concentration of the cells. Aliquots of 50 to 100 μL of concentrated *B. burgdorferi* cells in EPS were mixed with DNA and immediately electroporated in a 2 mm gap cuvette (BioRad) using the following settings: 2.5 kV, 25 μF, 200 Ω (122, 123). Shuttle vector DNA was used at 15-30 μg per transformation. Suicide vector DNA was linearized by restriction endonuclease digestion then ethanol-precipitated (124). Electroporated *B. burgdorferi* cells were resuspended in BSK-II medium, supplemented with 100 μM IPTG as needed, and allowed to recover until the next day. Transformants were selected by semisolid BSK-agarose plating under antibiotic selection as previously described (95). Single colonies were removed from the plates and expanded in BSK-II medium. Retention of the full parental plasmid complement was confirmed for each clone by multiplex PCR (125). Suicide vector transformants were checked for correct recombination by PCR. For shuttle vector transformants, 14 mL cultures were pelleted at 10,000 x g for 10 min, then the pellet was resuspended in 500 µL water and miniprepped using Zyppy plasmid miniprep kit (Zymo cat. D4020). The resulting DNA was used to transform NEB 5-alpha or NEB 5-alpha F’ lq *E. coli* competent cells (New England Biolabs). Colonies were selected using appropriate antibiotics, grown, miniprepped, and whole-plasmid sequenced using Nanopore technology at Quintara Biosciences.

### RNA isolation and qRT-PCR

*B. burgdorferi* cultures of strains VCbb07, VCbb08, and CNTs191 were maintained in exponential growth in the presence of 100 μM IPTG. The cells were pelleted by centrifugation for 10 min at 4,300 x g, resuspended in 1 mL IPTG-free BSK-II medium, their densities were determined by counting, and the cultures were diluted to 10^6^ cells/mL in 30 mL cultures with or without 100 μM IPTG. The next day, the cells were harvested by centrifugation (10 min at 4,300 x g, the pellet was resuspended in 400 μL buffer HN (126) and added to 1 mL RNAProtect Bacteria (Qiagen cat. 76506), vortexed, incubated for 5 min at room temperature, then centrifuged for 10 min at 10,000 x g. The liquid was removed by aspiration and the pellet was stored at -80°C. The RNA was released from the pellet using enzymatic lysis and proteinase K digestion performed as recommended by protocol 4 in the RNAprotect bacteria reagent manual. This was followed by purification using the RNeasy minikit (Qiagen cat. 74106). Trace DNA was removed using the Turbo DNA-free kit (Thermo Fisher Scientific cat. AM1907). The concentration of the RNA was determined using Nanodrop One^C^ and the RNA was stored at -80°C.

Quantitative reverse transcriptase PCR (qRT-PCR) was performed using the Verso 1-step RT-qPCR Kit (Thermo Scientific cat. AB4105C). The RNA was diluted to 10 ng/μL in water and 1 μL was used for each reaction. Two technical duplicate reactions were performed for each sample. An additional no RT control lacked the RT enzyme provided with the kit. The reaction volume was 25 μL/well. Cycling was done on a BioRad qRT-PCR system (CFX Duet) and the following conditions: reverse transcription, 15 min at 50°C; denaturation, 15 min at 95°C; 40 cycles each consisting of: 15 sec at 95°C, 30 sec at 50°C, 30 sec at 72°C with fluorescence reading in the SYBR channel; and a melting curve from 65°C to 95°C in 0.5°C increments. The primers used for qRT-PCR are listed in Table 4. Quantification cycle (C_q_) values and direct observation of amplification curves obtained in the absence of reverse transcription confirmed that the measured RNA amounts were not due to DNA contamination. Relative transcript amounts were determined by the ΔΔC_q_ method (127) and were normalized to the levels of the *flaB* transcript present in each sample.

**Table 4.**
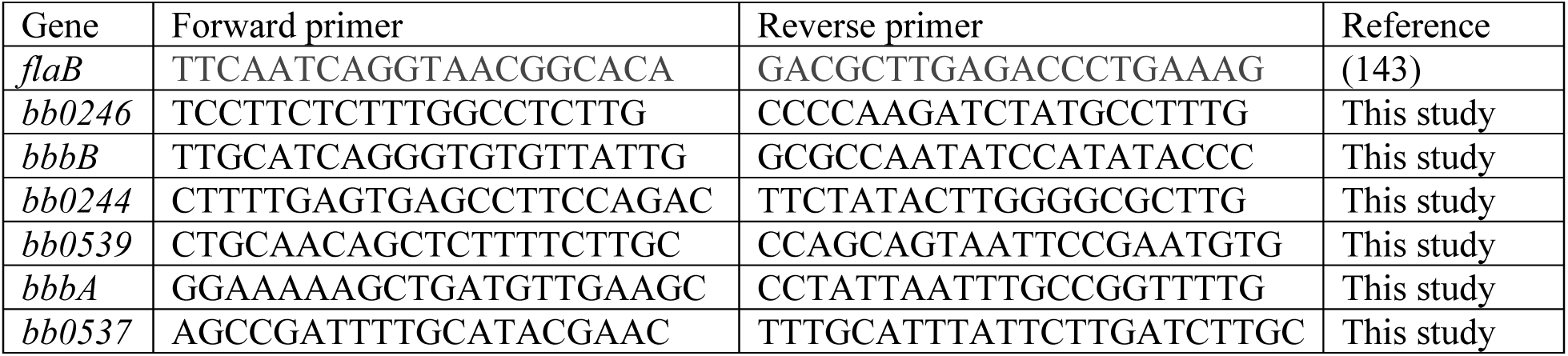
Primers used for qRT-PCR.

### RNA sequencing

RNA sequencing (RNA-seq) on *B. burgdorferi* strain B31-68-LS was performed previously (128). Briefly, total RNA was extracted from *B. burgdorferi* cells grown to culture densities between 5x10^7^ and 7x10^7^ cells/mL using TRIzol reagent (Thermo Fisher Scientific cat. 15596026) and treated with 1 unit of Ambion DNAse I (Thermo Fisher Scientific cat. AM2222) for 10 minutes at 37℃. RNA was quantified and assessed for quality, and RNA samples possessing RNA integrity number (RIN) values above 7.4 were used for sequencing. Ribosomal RNA was depleted using the QIAseq FastSelect -5S/16 S/23 S kit (Qiagen cat. 335925) before next generation sequencing library preparation with the TruSeq Stranded mRNA-Seq Sample Preparation Kit (Illumina cat. 20020595). Libraries were sequenced on an Illumina NextSeq 550 at 2 x 75 bp paired-end using a Mid Output 150 cycle kit (Illumina cat. 20024904). RNA-seq reads were compiled, filtered, and aligned to the *B. burgdorferi* B31 genome (RefSeq AE000783-AE000794 and AE001575-AE001584) using bowtie2 (129). Reads for annotated genes were quantified using featureCounts (130, 131). The RPKM (reads per kilobase per million reads) were then calculated to assess gene expression.

### Scanning electron microscopy

Strain VCbb08 was grown in culture medium supplemented with 100 µM IPTG or without IPTG supplementation for two days. Cells were fixed in 2% paraformaldehyde, 2.5% glutaraldehyde in 0.1 M Sorenson’s phosphate buffer for 30 min. Cells were then adhered to poly-L-lysine coated silicon chips (Electron Microscopy Sciences). Chips were processed as previously described (88): they were fixed in 0.5% OsO_4_, 0.8% K_4_Fe(CN)_6_ in 0.1 M sodium cacodylate buffer (Ted Pella cat. 18851) for 1 hour, rinsed in buffer, stained with 1% aqueous tannic acid for 1 hour, rinsed with buffer, and then one more fixation with 0.5% OsO_4_ + 0.8% K_4_Fe(CN)_6_ in buffer. Chips were then rinsed with distilled water and put through a graduated ethanol series into 100% ethanol. Chips were critically point dried using a Bal-Tec Critical Point Dryer 030 (Leica), mounted on stubs, and sputter coated with 10 nm of iridium (300T, Electron Microscopy Sciences). Images were acquired using a Hitachi SU8000 scanning electron microscope operating at 5 kV.

### Light microscopy

To collect phase contrast and fluorescence micrographs, *B. burgdorferi* cultures were spotted on a 2% agarose-PBS pad (93, 132) and then covered with a no. 1.5 coverslip. The cells were imaged using Nikon Ti inverted microscopes equipped with 100X Plan Apo 1.45 NA phase contrast oil objectives, Hamamatsu Orca-Flash4.0 V2 CMOS cameras and a Spectra X Light engine (Lumencor) or a Sola LE light source. The microscopes were equipped with the following Chroma filter cubes: DAPI (excitation ET395/25x, dichroic T425lpxr, emission ET460/50m); GFP (excitation ET470/40x, dichroic T495lpxr, emission ET525/50m); mCherry/TexasRed (excitation ET560/40x, dichroic T585lpxr, emission ET630/75m). Alternatively, images were acquired on a Nikon Ti2-E inverted microscope equipped with a 100X Plan Apo 1.45 NA phase contrast oil objective, a Hamamatsu Orca-Fusion BT camera, and a Spectra III LED fluorescent light source. The microscope was equipped with a custom filter cube containing a Semrock triple-edge polychroic (Semrock cat. FF409/493/596-Di02-t2-32x44-Ti2). Fluorescence images were captured with the excitation and emission filters: DAPI (excitation: 390/22, emission: 432/36), GFP (excitation: 475/28, emission: 525/50), and mCherry/TexasRed (excitation: 575/25, emission: 641/75). Images were acquired using NIS Elements software.

### Image analysis

Light microscopy images were analyzed as follows. First, cell outlines were generated based on the phase contrast images using the Oufti analysis package (133). Segmentation parameters were adjusted for each image set (Table S1) to optimize the automatically generated outlines. Outlines were curated, which involved: manual removal of outlines of cellular debris, of cells that curled on themselves or crossed other cells or of cells that were not fully within the image; manual extension of partial cell outlines or manual addition of full cell outlines, when feasible; manual splitting of cell outlines at sites of clear separation of cytoplasmic cylinders that remained linked by an outer membrane bridge (93). All outlines were refined using the “Refine All” function of the software, followed by a final removal of any improperly rendered outlines. Fluorescence background was subtracted before demographs and intensity profiles were generated, all using Oufti functions. Total cell fluorescence was calculated based on the final background subtracted and curated cell meshes using MATLAB (2024). The cell lists were then processed using the MATLAB script addMeshtoCellList.m and CalculateFluorPerCell.m as previously described (116). Final fluorescence data were plotted using the GraphPad Prism 10.0.0 software.

### Data visualization

The following software was used to generate, visualize, and present the data: Nikon NIS Elements software, FiJI (134), GraphPad Prism 10.0.0, Geneious Prime 2020.0.4, and Adobe Illustrator 2025.

### Labeling of new peptidoglycan insertion

New peptidoglycan insertion was labelled using the fluorescent D-amino acid HADA (Tocris Bioscience cat. 6647) following the procedure previously established in *B. burgdorferi* (93). *B. burgdorferi* cultures were grown under standard conditions described above. Cells were pelleted by centrifugation for 10 min at 4,300 x g, resuspended in 1 mL IPTG-free BSK-II medium, their densities were determined by counting, and the cultures were diluted to 10^6^ cells/mL in 6 mL cultures with or without 100 μM IPTG. After 24 hours growth, 990 µL of each culture was mixed with 10 µL 0.1 mM HADA in dimethyl sulfoxide (DMSO) to yield a final concentration of 1 µM HADA, and incubated for 1 hour at 34 °C. Following incubation, the cells were pelleted by centrifugation for 5 min 8,000 x g at 4 °C and resuspended by pipetting in 1 mL fridge-cold BSK-II with no added HADA or IPTG. Two additional washes were done, with the final pellet being resuspended in 30 – 100 µL fridge-cold BSK-II. Cells were imaged as described above.

### Phylogenetic analyses

Putative spirochete bactofilin protein sequences were identified using NCBI’s Basic Local Alignment Search Tool (BLAST) (135). The search was performed in June 2023. *Leptospira biflexa* bactofilin protein sequences LbbA through LbbE (NCBI RefSeq ID WP_012388301.1, WP_012390385.1, WP_012389582.1, WP_012388418.1, and WP_012388052.1) were used as bait and all hits within the spirochete phylum were retrieved. Additional searches were performed using *B. burgdorferi* strain B31 bactofilin hits (RefSeq ID WP_002665254.1, WP_002656070.1, and WP_002556844.1). The resulting spirochete bactofilin list was curated to remove duplicate entries. Only RefSeq entries (ID starting with WP) were retained. Additionally, partial sequences were removed. A total of 692 sequences were then assembled into a phylogenetic tree using Geneious Prime’s Geneious Tree Builder function and the following settings: alignment type, global alignment with free end gaps; cost matrix, Blosum62; genetic distance model, Jukes-Cantor; tree build method, neighbor-joining, with no outgroup. Sequence alignments were also performed using Geneious Prime’s Geneious Alignment function and the same alignment type and cost matrix.

### Structural predictions

Structural predictions were performed using the Alphafold3 platform (97, 136) hosted on Google’s Colaboratory platform (137). The structures predicted by Alphafold3 were visualized using UCSF ChimeraX 1.8 (138).

### Western blotting

Cultures of *B. burgdorferi* were grown as described in 45 mL liquid cultures in BSK-II. Conditions requiring IPTG depletion were created as described. Culture density was determined by direct counting and volumes were adjusted for a final input of 3.15 x 10^8^ cells. Cells were washed twice by pelleting (10,000 x g 10 min, 4 °C) and resuspension in 1 mL buffer HN containing 5X cOmplete protease inhibitor (Sigma Aldrich cat. 11836153001). Cell pellets were resuspended in 1X LDS sample buffer (Thermo Scientific cat. B0007) containing 5X cOmplete protease inhibitor, 1 mM DL-1,4-Dithiothreitol (Thermo Scientific cat. 426380100), and 2.5% (*v/v*) 2-mercaptoethanol then boiled at 95 °C for 5 min prior to loading for SDS-PAGE. Proteins were run on a 4-12% Bis-Tris Plus gel (Thermo Scientific cat. NW04122BOX) in 1X MES-SDS buffer (Thermo Scientific cat. NP0002) at 180 V for 25-30 min. Proteins were transferred to nitrocellulose membranes using an Invitrogen iBlot 2 gel transfer device (transfer settings: 20 V, 7 min). Membranes were washed with 0.1% (*v/v*) Tween-20 in 1X phosphate-buffered saline (PBS-T) and blocked with 4 % (*w/v*) bovine serum albumin in PBS-T. For mCherry detection, incubation with a primary anti-mCherry chicken polyclonal antibody (Novus Biologic cat. NBP2-25152) diluted 1:2,000 in blocking buffer was done overnight at 4 °C. Membranes were washed six times for 5 min each with PBS-T then incubated with a rabbit anti-chicken horseradish peroxidase (HRP)-conjugated secondary antibody (Invitrogen cat. A16130) diluted 1:2,000 in blocking buffer for 1 hour at room temperature. For FlaB, both primary and secondary incubations were done for 1 hour at room temperature with a rat anti-FlaB primary (Kerafast cat. ECN013) diluted 1:6,000 in blocking buffer, and rabbit anti-rat HRP-conjugated secondary (Sigma Aldrich cat. A5795) diluted 1:80,000. Chemiluminescence was detected after treatment with SuperSignal West PICO (Fisher Scientific cat. PI34577) or SuperSignal West FEMTO (Fisher Scientific cat. PI34094) PLUS chemiluminescent substrates using a BioRad Chemidoc gel imager.

## ACKNOWLEDGEMENTS

We are grateful to Dr. Scott D. Samuels from University of Montana for sharing plasmid pTA*flacp* and Dr. Jon Blevins from the University of Arkansas Medical School for sharing plasmid pJSB252 containing the P*_pQE30_* promoter.

We would like to thank the members of the Takacs laboratory for helpful discussions and feedback on the manuscript. V.C., C.L.S., J.W., P.E.S., and P.A.R. were supported by the Intramural Research Program of the National Institute of Allergy and Infectious Diseases, National Institutes of Health. Z.A.K. was supported in part by funding from the James D. Jamieson and Family M.D.-Ph.D. Scholarship Fund at Yale University and the Medical Scientist Training Grant T32 GM007205 from the National Institute of General Medical Sciences, National Institutes of Health. C.J.-W. is an investigator of the Howard Hughes Medical Institute. C.N.T. was supported in part by an American Heart Association postdoctoral fellowship (18POST33990330) and by startup funds from Northeastern University.

## AUTHOR CONTRIBUTIONS

Christopher B. Zinck, Conceptualization, Data Curation, Formal analysis, Investigation, Visualization, Writing – original draft, Writing – review and editing; Valentina Carracoi, Conceptualization, Investigation, Writing – review and editing; Zachary A. Kloos, Conceptualization, Investigation, Writing – review and editing; Cindi L. Schwartz, Investigation, Writing – review and editing; Jenny Wachter, Investigation, Writing – review and editing; Philip E. Stewart, Conceptualization, Supervision, Writing – review and editing; Christine Jacobs-Wagner, Conceptualization, Funding acquisition, Supervision, Writing – review and editing; Patricia A. Rosa, Conceptualization, Funding acquisition, Investigation, Formal analysis, Data Curation, Supervision, Writing – review and editing; Constantin N. Takacs, Conceptualization, Funding acquisition, Investigation, Data Curation, Formal analysis, Visualization, Supervision, Writing – original draft, Writing – review and editing.

## SUPPLEMENTARY FIGURE LEGENDS

**Figure S1.**
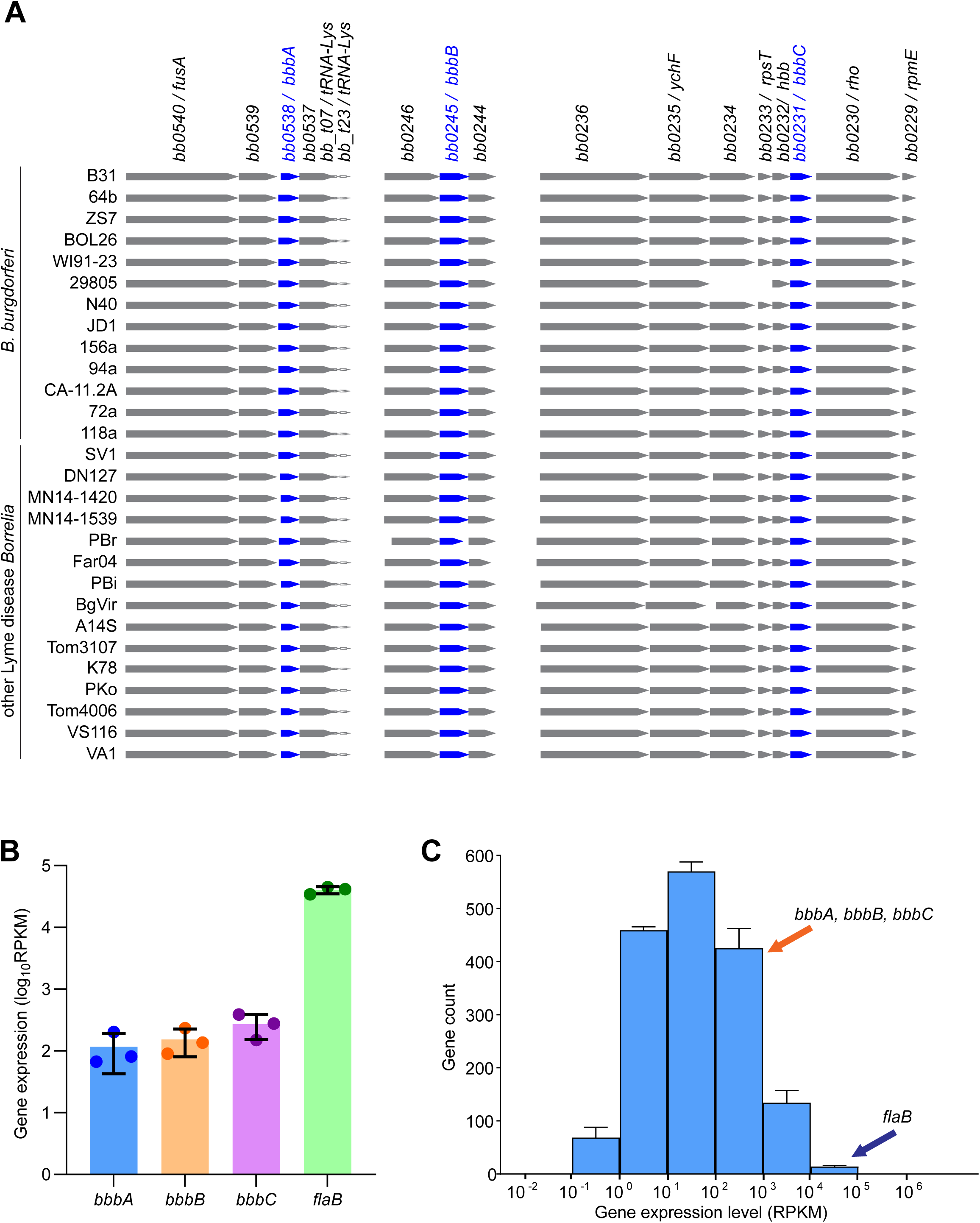
Genomic context and native expression of the *B. burgdorferi* bactofilins. **A.** Bactofilin loci in 29 Lyme disease *Borrelia* genomes visualized using the BorreliaBase online resource (96). The strains whose loci are depicted are listed on the left. Gene numbers listed at the top are based on the genome of the *B. burgdorferi* type strain B31. Bactofilin-encoding genes are shown in blue. **B.** Bactofilin gene expression detected by RNA-seq in strain B31-68-LS during in vitro cultivation. The highly expressed *flaB* gene is included for reference. Shown are individual values as well as means ± standard deviations of *n*=3 replicates. The RNA-seq data was published in a previous study (128). RPKM, reads per kilobase per million reads. **C.** Histogram depicting the expression of all the genes detected in strain B31-68-LS during in vitro cultivation. The genes are grouped in successive 10-fold expression bins from 10^-2^ to 10^6^ RPKM. Each bar depicts the mean number of genes that belong to each expression bin, while the error bars depict the standard deviation of this number for the *n*=3 replicates analyzed. The bins containing the three bactofilin genes *bbbA*, *bbbB*, and *bbbC* are indicated by orange arrows. For reference, the bin containing the highly expressed *flaB* gene is indicated by a blue arrow.

**Figure S2.**
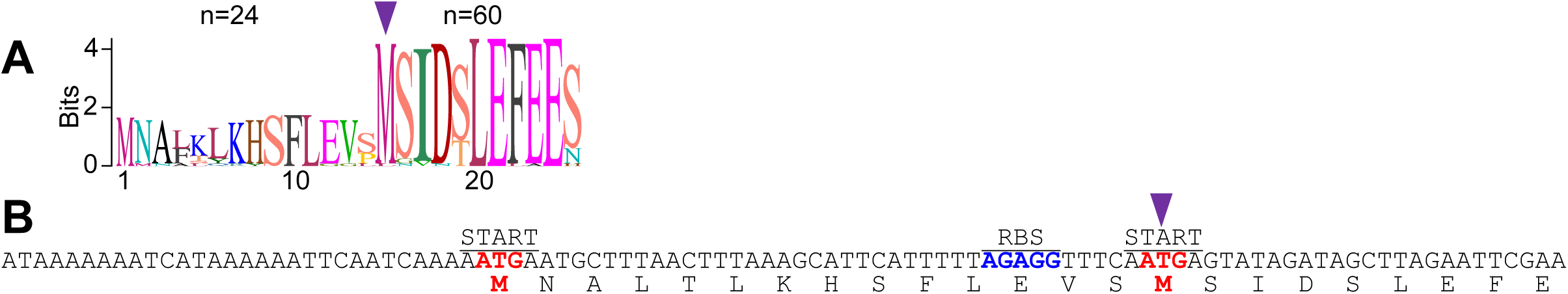
Evidence for likely bioinformatic mis-annotation of a subset of BbbA sequences. **A.** Sequence logo of 60 *B. burgdorferi* BbbA sequences. Some of these sequences (*n*=24) have an N-terminal tail extension of 14 amino acids. The 15^th^ amino acid (indicated by an arrowhead) is a conserved START codon in all 60 sequences (*n*=59 sequences have a methionine residue while one sequence has a lysine residue at this position, which can function as a start codon in bacteria). **B.** Nucleotide sequence of the *B. burgdorferi* B31 genome (139) near the beginning of the *bbbA* gene and its encoded protein sequence. Nucleotide bases are listed from left to right in the 5’ to 3’ direction of the coding strand of *bbbA*. Putative start codons are in red, with the leftmost being the annotated start codon. Encoded amino acids are listed below the nucleotide sequence using the single letter convention. Likely ribosomal binding site (RBS) is in blue. Functional start codon used in this study is indicated by the same arrowhead as in (A).

**Figure S3.**
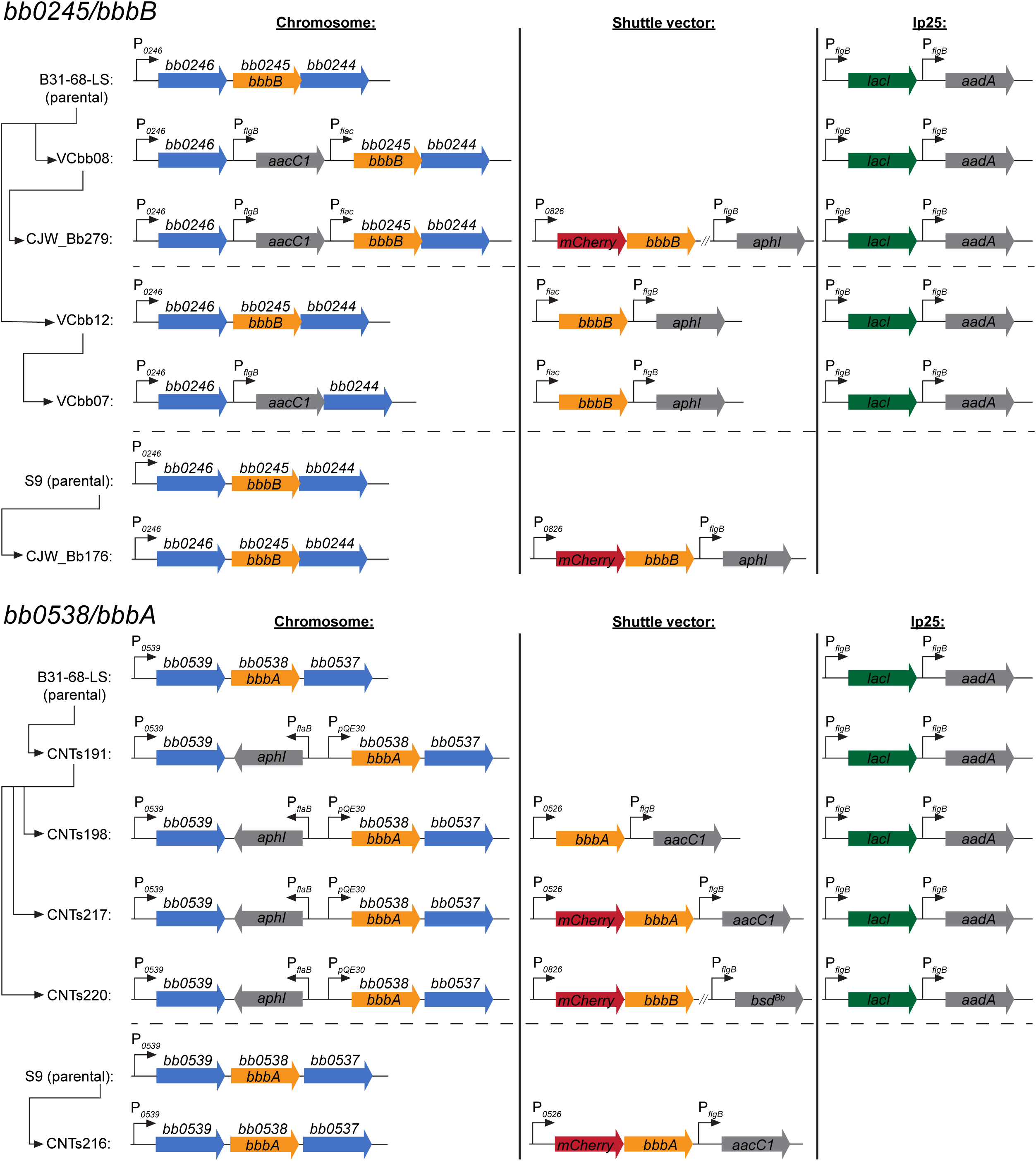
Schematic depictions of relevant gene features of the strains used in this study. Relevant features of strains generated to investigate bactofilin BbbB (top) and BbbA (bottom). The chromosomal loci are depicted in the left column. If present, relevant genes expressed from the shuttle vector are listed in the middle column. For strains derived from B31-68-LS, LacIBb expression from endogenous plasmid lp25 is shown in the right column. Arrows at the left identify the strain lineages, which are separated using horizontal lines.

**Figure S4.**
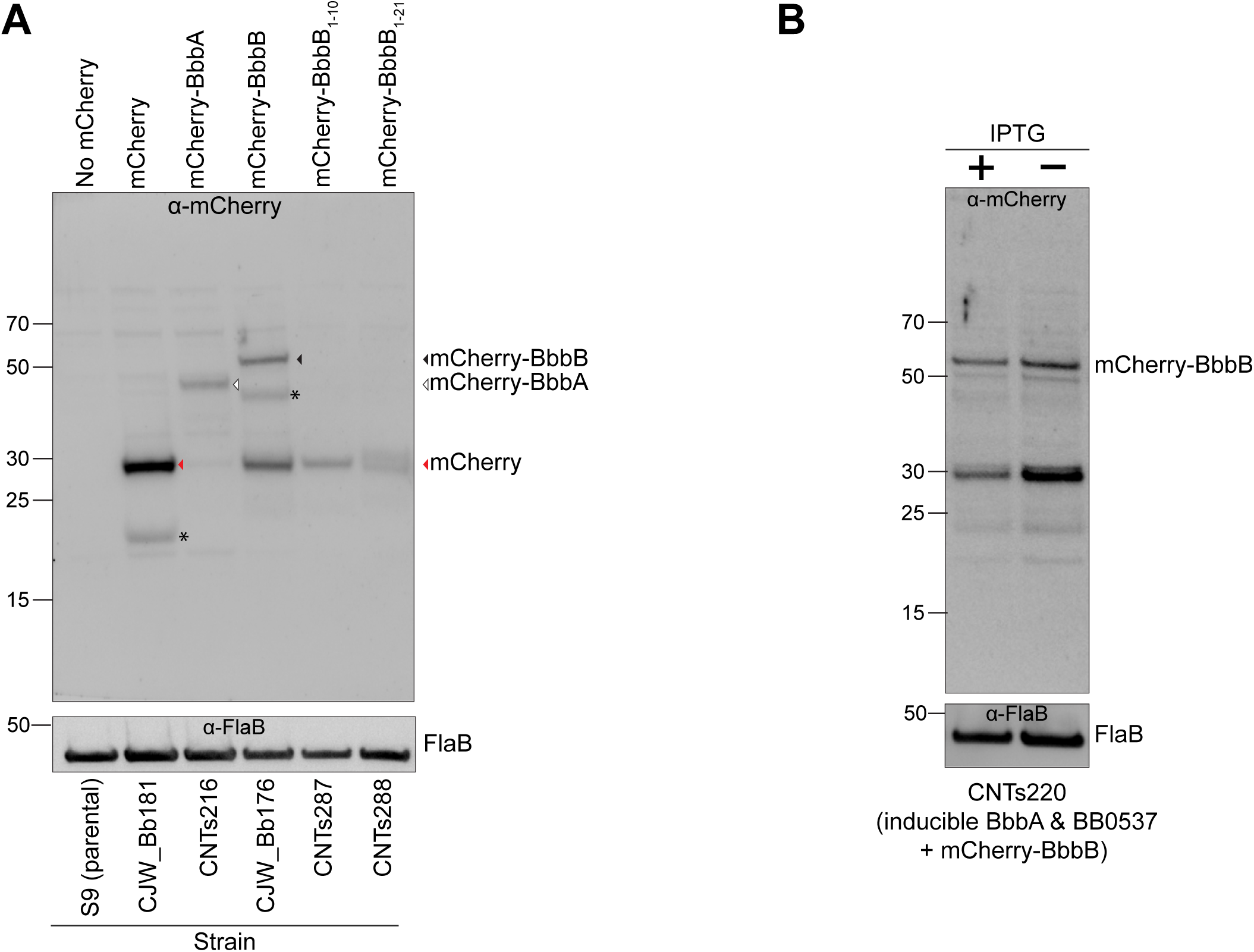
Western blotting detection of mCherry-bactofilin fusions. **A.** Western blotting of lysates of strains used in this study. Top, blotting with an anti-mCherry antibody. Bottom, blotting with anti-FlaB antibody of the same membrane used for the anti-mCherry blot. Identities of the major bands are given at the right, indicated by corresponding arrows. Protein size standards (kDa) are indicated at the left. Strain identities are at the bottom. Relevant expressed proteins are at the top. Bands marked by an asterisk are consistent with mCherry degradation products generated during protein boiling prior to SDS-PAGE (140). **B.** Western blotting of lysates of strain CNTs220 (carrying IPTG-inducible *bbbA* and *bb0537* along with shuttle vector-borne, constitutively expressed *mcherry-bbbB*) grown with 1 mM IPTG or grown in the absence of IPTG for 24 h. Top, blotting with an anti-mCherry antibody. Bottom, blotting with anti-FlaB antibody of the same membrane used for the anti-mCherry blot.

**Figure S5.**
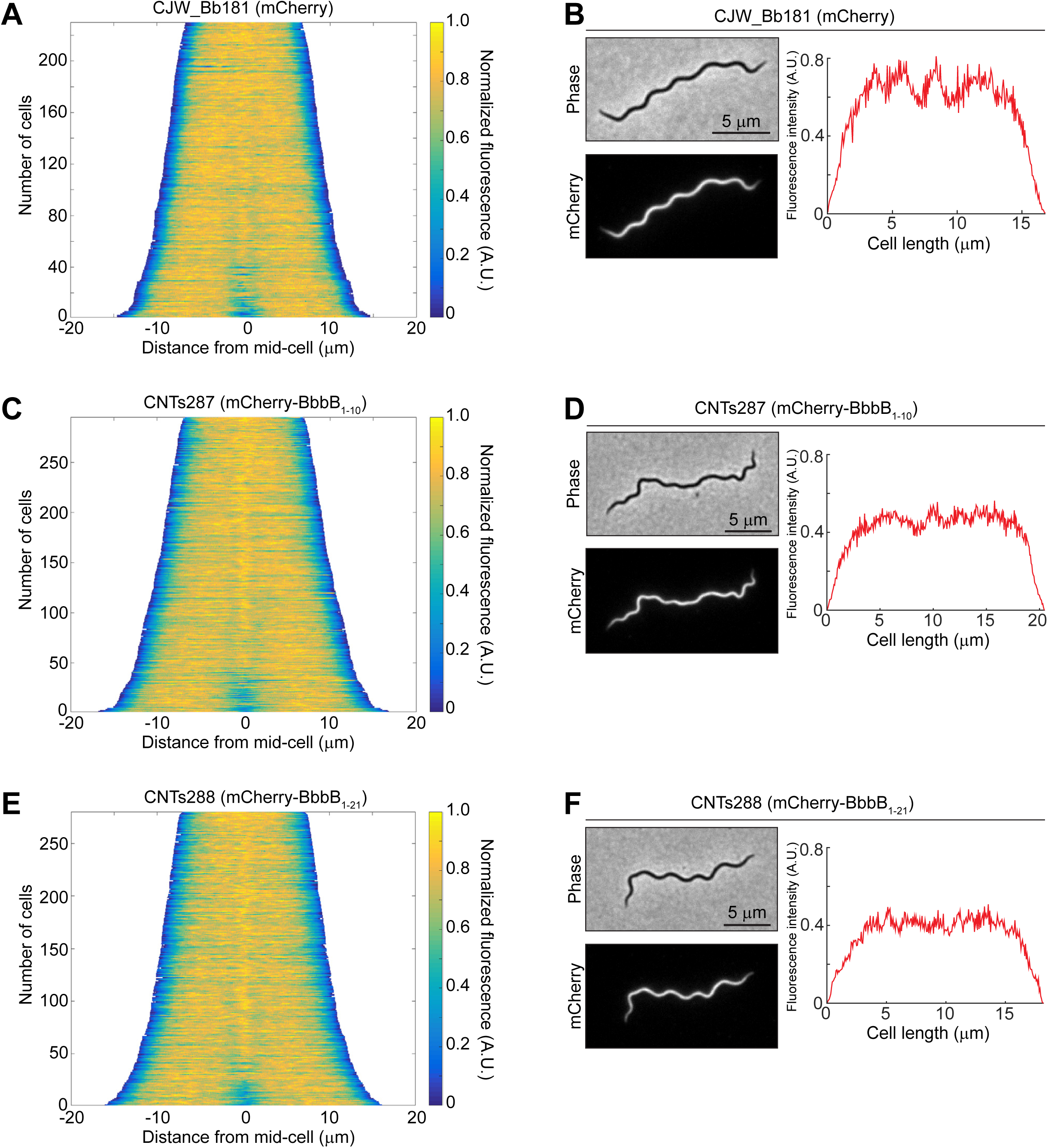
Localization of free mCherry and mCherry fused to BbbB N-terminal fragments. A. Demograph of a population of 230 cells of strain CJW_Bb181 expressing free mCherry. A.U., arbitrary units. B. Phase contrast and fluorescence micrographs (left), and line intensity profile (right) of a cell of strain CJW_Bb181 expressing free mCherry. A.U., arbitrary units. C. Same as in A but for a population of 293 cells of strain CNTs287 expressing mCherry-BbbB_1-10_. D. Same as in B but for a cell of strain CNTs287. E. Same as in A but for a population of 280 cells of strain CNTs288 expressing mCherry-BbbB_1-21_. F. Same as in B but for a cell of strain CNTs288.

**Figure S6.**
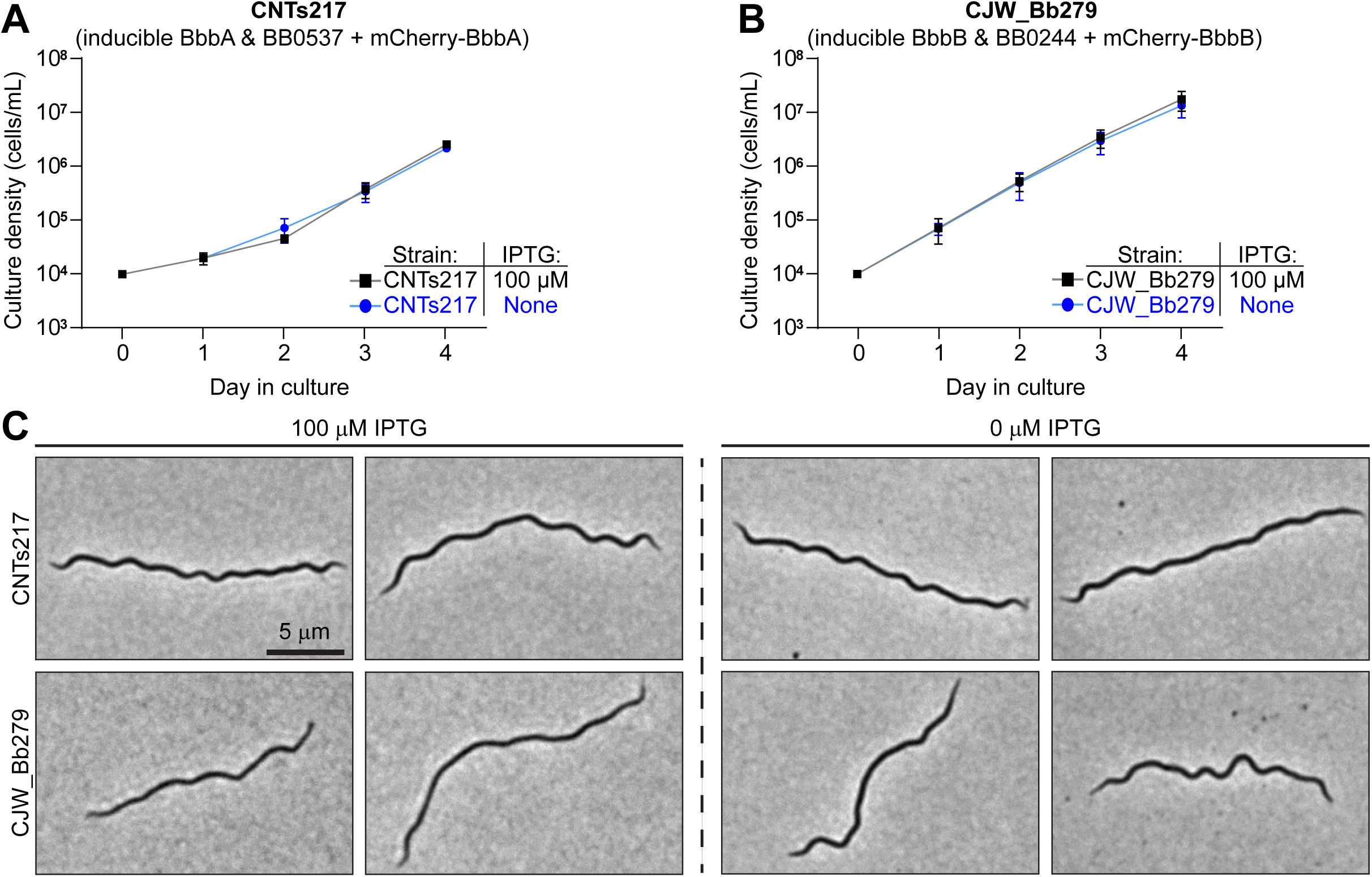
Complementation of bactofilin depletion by mCherry-bactofilin fusions. **A.** Growth curve of strain CNTs217 (carrying IPTG-inducible *bbbA* and *bb0537* along with shuttle vector-borne constitutively expressed *mcherry-bbbA*) grown in the presence or absence of 100 µM IPTG. Points are means ± standard deviations of results from *n*=3 cultures. **B.** Growth curve of strain CJW_Bb279 (carrying IPTG-inducible *bbbB* and *bb0244* along with shuttle-vector borne constitutively expressed *mcherry-bbbB*) grown in the presence or absence of 100 µM IPTG. Points are means ± standard deviations of results from *n*=3 cultures. **C.** Phase contrast light micrographs of cells of strains CNTs217 (top) and CJW_Bb279 (bottom) grown with 100 μM IPTG (left) or for 48 h without IPTG (0 μM IPTG, right). Two images are provided for each condition.

**Figure S7.**
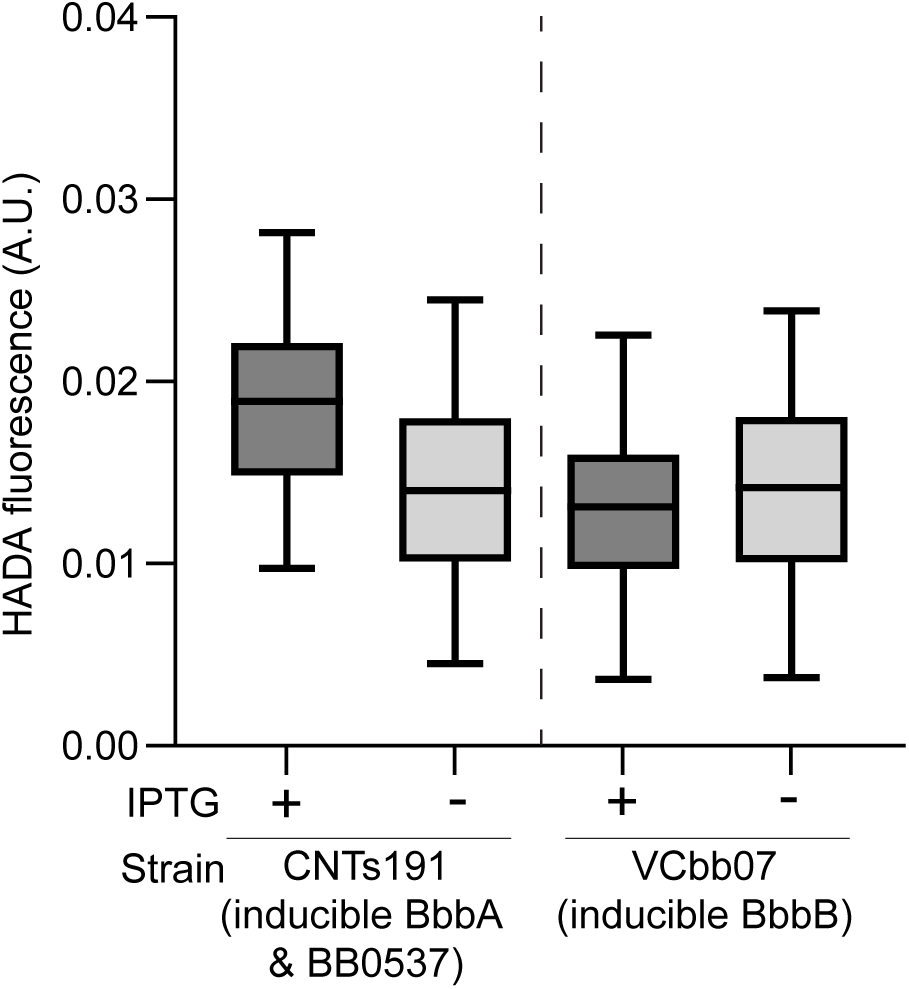
Quantification of HADA uptake by bactofilin depletion strains. Total HADA fluorescence normalized by cell area for populations of cells of strain CNTs191 (carrying IPTG-inducible *bbbA* and *bb0537*, left) or strain VCbb07 (carrying IPTG-inducible *bbbB*, right) with or without IPTG induction. Cell populations are the same used in the demographs in Figs. 8A, 8C, 8G, and 8I, respectively). Boxes represent the interquartile ranges of cell fluorescence with the mean as the midline; tails represent the 2.5 and 97.5 percentiles.

## REFERENCES

1. Radolf JD, Caimano MJ, Stevenson B, Hu LT. 2012. Of ticks, mice and men: understanding the dual-host lifestyle of Lyme disease spirochaetes. Nature Reviews Microbiology 10:87–99.

2. Bourgeois JS, Hu LT. 2024. Hitchhiker’s Guide to *Borrelia burgdorferi*. J Bacteriol 206:e0011624.

3. Steere AC, Strle F, Wormser GP, Hu LT, Branda JA, Hovius JWR, Li X, Mead PS. 2016. Lyme borreliosis. Nature Reviews Disease Primers 2:16090.

4. Mead PS. 2015. Epidemiology of Lyme disease. Infect Dis Clin North Am 29:187–210.

5. Kugeler KJ, Earley A, Mead PS, Hinckley AF. 2024. Surveillance for Lyme Disease After Implementation of a Revised Case Definition - United States, 2022. MMWR Morb Mortal Wkly Rep 73:118–123.

6. Pustijanac E, Bursic M, Millotti G, Paliaga P, Ivesa N, Cvek M. 2024. Tick-borne bacterial diseases in Europe: Threats to public health. Eur J Clin Microbiol Infect Dis 43:1261–1295.

7. De Silva AM, Fikrig E. 1995. Growth and migration of *Borrelia burgdorferi* in *Ixodes* ticks during blood feeding. Am J Trop Med Hyg 53:397–404.

8. Piesman J, Oliver JR, Sinsky RJ. 1990. Growth kinetics of the Lyme disease spirochete (*Borrelia burgdorferi*) in vector ticks (*Ixodes dammini*). Am J Trop Med Hyg 42:352–7.

9. Piesman J, Sinsky RJ. 1988. Ability of *Ixodes scapularis*, *Dermacentor variabilis*, and *Amblyomma americanum* (Acari: Ixodidae) to acquire, maintain, and transmit Lyme disease spirochetes (*Borrelia burgdorferi*). J Med Entomol 25:336-9.

10. Piesman J, Mather TN, Sinsky RJ, Spielman A. 1987. Duration of tick attachment and *Borrelia burgdorferi* transmission. J Clin Microbiol 25:557–8.

11. Hyde JA. 2017. *Borrelia burgdorferi* keeps moving and carries on: A review of Borrelial dissemination and invasion. Front Immunol 8:114.

12. Casselli T, Divan A, Vomhof-DeKrey EE, Tourand Y, Pecoraro HL, Brissette CA. 2021. A murine model of Lyme disease demonstrates that *Borrelia burgdorferi* colonizes the dura mater and induces inflammation in the central nervous system. PLoS Pathog 17:e1009256.

13. Barthold SW, Beck DS, Hansen GM, Terwilliger GA, Moody KD. 1990. Lyme borreliosis in selected strains and ages of laboratory mice. J Infect Dis 162:133–8.

14. Barthold SW, Persing DH, Armstrong AL, Peeples RA. 1991. Kinetics of *Borrelia burgdorferi* dissemination and evolution of disease after intradermal inoculation of mice. Am J Pathol 139:263–73.

15. Barthold SW, de Souza MS, Janotka JL, Smith AL, Persing DH. 1993. Chronic Lyme borreliosis in the laboratory mouse. Am J Pathol 143:959–71.

16. Yang L, Weis JH, Eichwald E, Kolbert CP, Persing DH, Weis JJ. 1994. Heritable susceptibility to severe *Borrelia burgdorferi*-induced arthritis is dominant and is associated with persistence of large numbers of spirochetes in tissues. Infect Immun 62:492–500.

17. Muehlenbachs A, Bollweg BC, Schulz TJ, Forrester JD, DeLeon Carnes M, Molins C, Ray GS, Cummings PM, Ritter JM, Blau DM, Andrew TA, Prial M, Ng DL, Prahlow JA, Sanders JH, Shieh WJ, Paddock CD, Schriefer ME, Mead P, Zaki SR. 2016. Cardiac tropism of *Borrelia burgdorfer*i: An autopsy study of sudden cardiac death associated with Lyme carditis. Am J Pathol 186:1195–205.

18. Coburn J, Garcia B, Hu LT, Jewett MW, Kraiczy P, Norris SJ, Skare J. 2021. Lyme Disease pathogenesis. Curr Issues Mol Biol 42:473–518.

19. Steere AC. 2024. Lyme Arthritis: A 50-year journey. J Infect Dis 230:S1–S10.

20. Baarsma ME, Hovius JW. 2024. Persistent symptoms after Lyme disease: Clinical characteristics, predictors, and classification. J Infect Dis 230:S62–S69.

21. Cardenas-de la Garza JA, De la Cruz-Valadez E, Ocampo-Candiani J, Welsh O. 2019. Clinical spectrum of Lyme disease. Eur J Clin Microbiol Infect Dis 38:201–208.

22. Verschoor YL, Vrijlandt A, Spijker R, van Hest RM, Ter Hofstede H, van Kempen K, Henningsson AJ, Hovius JW. 2022. Persistent *Borrelia burgdorferi* sensu lato infection after antibiotic treatment: Systematic overview and appraisal of the current evidence from experimental animal models. Clin Microbiol Rev 35:e0007422.

23. Zafar K, Azuama OC, Parveen N. 2024. Current and emerging approaches for eliminating *Borrelia burgdorferi* and alleviating persistent Lyme disease symptoms. Front Microbiol 15:1459202.

24. Lantos PM, Rumbaugh J, Bockenstedt LK, Falck-Ytter YT, Aguero-Rosenfeld ME, Auwaerter PG, Baldwin K, Bannuru RR, Belani KK, Bowie WR, Branda JA, Clifford DB, DiMario FJ, Halperin JJ, Krause PJ, Lavergne V, Liang MH, Meissner HC, Nigrovic LE, Nocton JJJ, Osani MC, Pruitt AA, Rips J, Rosenfeld LE, Savoy ML, Sood SK, Steere AC, Strle F, Sundel R, Tsao J, Vaysbrot EE, Wormser GP, Zemel LS. 2021. Clinical practice guidelines by the Infectious Diseases Society of America (IDSA), American Academy of Neurology (AAN), and American College of Rheumatology (ACR): 2020 guidelines for the prevention, diagnosis and treatment of Lyme disease. Clin Infect Dis 72:e1–e48.

25. Aucott JN, Yang T, Yoon I, Powell D, Geller SA, Rebman AW. 2022. Risk of post-treatment Lyme disease in patients with ideally-treated early Lyme disease: A prospective cohort study. Int J Infect Dis 116:230–237.

26. Weitzner E, McKenna D, Nowakowski J, Scavarda C, Dornbush R, Bittker S, Cooper D, Nadelman RB, Visintainer P, Schwartz I, Wormser GP. 2015. Long-term assessment of post-treatment symptoms in patients with culture-confirmed early Lyme disease. Clin Infect Dis 61:1800–6.

27. Marques A. 2022. Persistent symptoms after treatment of Lyme disease. Infect Dis Clin North Am 36:621–638.

28. Dunham-Ems SM, Caimano MJ, Pal U, Wolgemuth CW, Eggers CH, Balic A, Radolf JD. 2009. Live imaging reveals a biphasic mode of dissemination of *Borrelia burgdorferi* within ticks. J Clin Invest 119:3652–65.

29. Zhang J, Takacs CN, McCausland JW, Mueller EA, Buron J, Thappeta Y, Wachter J, Rosa PA, Jacobs-Wagner C. 2025. *Borrelia burgdorferi* loses essential genetic elements and cell proliferative potential during stationary phase in culture but not in the tick vector. J Bacteriol doi:10.1128/jb.00457-24:e0045724.

30. Koloski CW, Hurry G, Foley-Eby A, Adam H, Goldstein S, Zvionow P, Detmer SE, Voordouw MJ. 2024. Male C57BL/6J mice have higher presence and abundance of *Borrelia burgdorferi* in their ventral skin compared to female mice. Ticks Tick Borne Dis 15:102308.

31. Rego RO, Bestor A, Stefka J, Rosa PA. 2014. Population bottlenecks during the infectious cycle of the Lyme disease spirochete *Borrelia burgdorferi*. PLoS One 9:e101009.

32. Norris SJ. 2014. Antigenic variation systems of Lyme disease *Borrelia*: Eluding host immunity through both random, segmental gene conversion and framework heterogeneity. Microbiology Spectrum 2.

33. Chaconas G, Castellanos M, Verhey TB. 2020. Changing of the guard: How the Lyme disease spirochete subverts the host immune response. J Biol Chem 295:301–313.

34. Crowder CD, Denny RL, Barbour AG. 2017. Segregation lag in polyploid cells of the pathogen genus *Borrelia*: Implications for antigenic variation. Yale Journal of Biology and Medicine 90:195–218.

35. Aranjuez GF, Kuhn HW, Adams PP, Jewett MW. 2019. *Borrelia burgdorferi bbk13* is critical for spirochete population expansion in the skin during early infection. Infect Immun 87:e00887–18.

36. Aranjuez GF, Lasseter AG, Jewett MW. 2021. The infectivity gene *bbk13* is important for multiple phases of the *Borrelia burgdorferi* enzootic cycle. Infect Immun 89:e0021621.

37. Egan AJF, Errington J, Vollmer W. 2020. Regulation of peptidoglycan synthesis and remodelling. Nat Rev Microbiol 18:446–460.

38. Rohs PDA, Bernhardt TG. 2021. Growth and division of the peptidoglycan matrix. Annu Rev Microbiol 75:315–336.

39. Holtje JV. 1998. Growth of the stress-bearing and shape-maintaining murein sacculus of *Escherichia coli*. Microbiol Mol Biol Rev 62:181–203.

40. Wolgemuth CW, Charon NW, Goldstein SF, Goldstein RE. 2006. The flagellar cytoskeleton of the spirochetes. J Mol Microbiol Biotechnol 11:221–7.

41. Raddi G, Morado DR, Yan J, Haake DA, Yang XF, Liu J. 2012. Three-dimensional structures of pathogenic and saprophytic *Leptospira* species revealed by cryo-electron tomography. J Bacteriol 194:1299–306.

42. Kudryashev M, Cyrklaff M, Baumeister W, Simon MM, Wallich R, Frischknecht F. 2009. Comparative cryo-electron tomography of pathogenic Lyme disease spirochetes. Mol Microbiol 71:1415–34.

43. Caimano MJ, Bourell KW, Bannister TD, Cox DL, Radolf JD. 1999. The *Treponema denticola* major sheath protein is predominantly periplasmic and has only limited surface exposure. Infect Immun 67:4072–83.

44. Izard J, Renken C, Hsieh CE, Desrosiers DC, Dunham-Ems S, La Vake C, Gebhardt LL, Limberger RJ, Cox DL, Marko M, Radolf JD. 2009. Cryo-electron tomography elucidates the molecular architecture of *Treponema pallidum*, the syphilis spirochete. J Bacteriol 191:7566–80.

45. Fontana C, Lambert A, Benaroudj N, Gasparini D, Gorgette O, Cachet N, Bomchil N, Picardeau M. 2016. Analysis of a spontaneous non-motile and avirulent mutant shows that FliM is required for full endoflagella assembly in *Leptospira interrogans*. PLoS One 11:e0152916.

46. Charon NW, Cockburn A, Li C, Liu J, Miller KA, Miller MR, Motaleb MA, Wolgemuth CW. 2012. The unique paradigm of spirochete motility and chemotaxis. Annu Rev Microbiol 66:349–70.

47. Ruby JD, Li H, Kuramitsu H, Norris SJ, Goldstein SF, Buttle KF, Charon NW. 1997. Relationship of *Treponema denticola* periplasmic flagella to irregular cell morphology. J Bacteriol 179:1628–35.

48. Rosey EL, Kennedy MJ, Petrella DK, Ulrich RG, Yancey RJ, Jr. 1995. Inactivation of *Serpulina hyodysenteriae flaA1* and *flaB1* periplasmic flagellar genes by electroporation-mediated allelic exchange. J Bacteriol 177:5959–70.

49. Holt SC. 1978. Anatomy and chemistry of spirochetes. Microbiol Rev 42:114–60.

50. Motaleb MA, Corum L, Bono JL, Elias AF, Rosa P, Samuels DS, Charon NW. 2000. *Borrelia burgdorferi* periplasmic flagella have both skeletal and motility functions. Proc Natl Acad Sci U S A 97:10899–904.

51. Attaibi M, den Blaauwen T. 2022. An updated model of the divisome: Regulation of the septal peptidoglycan synthesis machinery by the divisome. Int J Mol Sci 23:3537.

52. Morrison JJ, Camberg JL. 2024. Building the bacterial divisome at the septum, p 49–71, Subcellular Biochemistry doi:10.1007/978-3-031-58843-3_4. Springer International Publishing.

53. Shi H, Bratton BP, Gitai Z, Huang KC. 2018. How to build a bacterial cell: MreB as the foreman of *E. coli* construction. Cell 172:1294–1305.

54. Cameron TA, Zupan JR, Zambryski PC. 2015. The essential features and modes of bacterial polar growth. Trends Microbiol 23:347–53.

55. Flardh K. 2003. Essential role of DivIVA in polar growth and morphogenesis in *Streptomyces coelicolor* A3(2). Mol Microbiol 49:1523–36.

56. Kang CM, Nyayapathy S, Lee JY, Suh JW, Husson RN. 2008. Wag31, a homologue of the cell division protein DivIVA, regulates growth, morphology and polar cell wall synthesis in mycobacteria. Microbiology 154:725–735.

57. Letek M, Ordonez E, Vaquera J, Margolin W, Flardh K, Mateos LM, Gil JA. 2008. DivIVA is required for polar growth in the MreB-lacking rod-shaped actinomycete *Corynebacterium glutamicum*. J Bacteriol 190:3283–92.

58. Brown PJ, de Pedro MA, Kysela DT, Van der Henst C, Kim J, De Bolle X, Fuqua C, Brun YV. 2012. Polar growth in the Alphaproteobacterial order Rhizobiales. Proc Natl Acad Sci U S A 109:1697–701.

59. Umeda A, Amako K. 1983. Growth of the surface of *Corynebacterium diphtheriae*. Microbiol Immunol 27:663–71.

60. Thanky NR, Young DB, Robertson BD. 2007. Unusual features of the cell cycle in mycobacteria: polar-restricted growth and the snapping-model of cell division. Tuberculosis (Edinb) 87:231–6.

61. Garde S, Chodisetti PK, Reddy M. 2021. Peptidoglycan: Structure, synthesis, and regulation. EcoSal Plus 9:ecosalplus.ESP-0010-2020.

62. Galinier A, Delan-Forino C, Foulquier E, Lakhal H, Pompeo F. 2023. Recent advances in peptidoglycan synthesis and regulation in bacteria. Biomolecules 13:720.

63. van Teeffelen S, Wang S, Furchtgott L, Huang KC, Wingreen NS, Shaevitz JW, Gitai Z. 2011. The bacterial actin MreB rotates, and rotation depends on cell-wall assembly. Proc Natl Acad Sci U S A 108:15822–7.

64. Dominguez-Escobar J, Chastanet A, Crevenna AH, Fromion V, Wedlich-Soldner R, Carballido-Lopez R. 2011. Processive movement of MreB-associated cell wall biosynthetic complexes in bacteria. Science 333:225–8.

65. Garner EC, Bernard R, Wang W, Zhuang X, Rudner DZ, Mitchison T. 2011. Coupled, circumferential motions of the cell wall synthesis machinery and MreB filaments in *B. subtilis*. Science 333:222–225.

66. Kelemen GH. 2017. Intermediate filaments supporting cell shape and growth in bacteria, p 161–211, Subcellular Biochemistry doi:10.1007/978-3-319-53047-5_6. Springer International Publishing.

67. Ausmees N, Kuhn JR, Jacobs-Wagner C. 2003. The bacterial cytoskeleton: an intermediate filament-like function in cell shape. Cell 115:705–13.

68. Cabeen MT, Charbon G, Vollmer W, Born P, Ausmees N, Weibel DB, Jacobs-Wagner C. 2009. Bacterial cell curvature through mechanical control of cell growth. EMBO J 28:1208–19.

69. Cabeen MT, Murolo MA, Briegel A, Bui NK, Vollmer W, Ausmees N, Jensen GJ, Jacobs-Wagner C. 2010. Mutations in the lipopolysaccharide biosynthesis pathway interfere with crescentin-mediated cell curvature in *Caulobacter crescentus*. J Bacteriol 192:3368–78.

70. Deng X, Gonzalez Llamazares A, Wagstaff JM, Hale VL, Cannone G, McLaughlin SH, Kureisaite-Ciziene D, Lowe J. 2019. The structure of bactofilin filaments reveals their mode of membrane binding and lack of polarity. Nat Microbiol 4:2357–2368.

71. Curtis Z, Escudeiro P, Mallon J, Leland O, Rados T, Dodge A, Andre K, Kwak J, Yun K, Isaac B, Martinez Pastor M, Schmid AK, Pohlschroder M, Alva V, Bisson A. 2024. Halofilins as emerging bactofilin families of archaeal cell shape plasticity orchestrators. Proc Natl Acad Sci U S A 121:e2401583121.

72. Kuhn J, Briegel A, Morschel E, Kahnt J, Leser K, Wick S, Jensen GJ, Thanbichler M. 2010. Bactofilins, a ubiquitous class of cytoskeletal proteins mediating polar localization of a cell wall synthase in *Caulobacter crescentus*. EMBO J 29:327–39.

73. Shi C, Fricke P, Lin L, Chevelkov V, Wegstroth M, Giller K, Becker S, Thanbichler M, Lange A. 2015. Atomic-resolution structure of cytoskeletal bactofilin by solid-state NMR. Sci Adv 1:e1501087.

74. Lee J, Cox JV, Ouellette SP. 2023. The unique N-Terminal domain of Chlamydial bactofilin mediates its membrane localization and ring-forming properties. J Bacteriol 205:e0009223.

75. Zuckerman DM, Boucher LE, Xie K, Engelhardt H, Bosch J, Hoiczyk E. 2015. The bactofilin cytoskeleton protein BacM of *Myxococcus xanthus* forms an extended beta-sheet structure likely mediated by hydrophobic interactions. PLoS One 10:e0121074.

76. Caccamo PD, Jacq M, VanNieuwenhze MS, Brun YV. 2020. A division of labor in the recruitment and topological organization of a bacterial morphogenic complex. Curr Biol 30:3908–3922 e4.

77. Liu Y, Karmakar R, Steinchen W, Mukherjee S, Bange G, Schäfer LV, Thanbichler M. 2024. Membrane binding properties of the cytoskeletal protein bactofilin. doi:10.1101/2024.06.14.599034.

78. Holtrup S, Graumann PL. 2022. Strain-dependent motility defects and suppression by a *flhO* mutation for *B. subtilis* bactofilins. BMC Res Notes 15:168.

79. Lin L, Osorio Valeriano M, Harms A, Sogaard-Andersen L, Thanbichler M. 2017. Bactofilin-mediated organization of the ParABS chromosome segregation system in *Myxococcus xanthus*. Nat Commun 8:1817.

80. Anand D, Schumacher D, Sogaard-Andersen L. 2020. SMC and the bactofilin/PadC scaffold have distinct yet redundant functions in chromosome segregation and organization in *Myxococcus xanthus*. Mol Microbiol 114:839–856.

81. Bulyha I, Lindow S, Lin L, Bolte K, Wuichet K, Kahnt J, van der Does C, Thanbichler M, Sogaard-Andersen L. 2013. Two small GTPases act in concert with the bactofilin cytoskeleton to regulate dynamic bacterial cell polarity. Dev Cell 25:119–31.

82. Brockett MR, Lee J, Cox JV, Liechti GW, Ouellette SP. 2021. A dynamic, ring-forming bactofilin critical for maintaining cell size in the obligate intracellular bacterium *Chlamydia trachomatis*. Infect Immun 89:e0020321.

83. Richter P, Melzer B, Muller FD. 2023. Interacting bactofilins impact cell shape of the MreB-less multicellular *Rhodomicrobium vannielii*. PLoS Genet 19:e1010788.

84. Pöhl S, Osorio-Valeriano M, Cserti E, Harberding J, Hernandez-Tamayo R, Biboy J, Sobetzko P, Vollmer W, Graumann PL, Thanbichler M. 2024. A dynamic bactofilin cytoskeleton cooperates with an M23 endopeptidase to control bacterial morphogenesis. eLife 12:RP86577.

85. Holtrup S, Heimerl T, Linne U, Altegoer F, Noll F, Waidner B. 2019. Biochemical characterization of the *Helicobacter pylori* bactofilin-homolog HP1542. PLoS One 14:e0218474.

86. Frirdich E, Vermeulen J, Biboy J, Vollmer W, Gaynor EC. 2023. Multiple *Campylobacter jejuni* proteins affecting the peptidoglycan structure and the degree of helical cell curvature. Front Microbiol 14:1162806.

87. Sycuro LK, Pincus Z, Gutierrez KD, Biboy J, Stern CA, Vollmer W, Salama NR. 2010. Peptidoglycan crosslinking relaxation promotes *Helicobacter pylori*’s helical shape and stomach colonization. Cell 141:822–33.

88. Jackson KM, Schwartz C, Wachter J, Rosa PA, Stewart PE. 2018. A widely conserved bacterial cytoskeletal component influences unique helical shape and motility of the spirochete *Leptospira biflexa*. Mol Microbiol 108:77–89.

89. Razew A, Schwarz JN, Mitkowski P, Sabala I, Kaus-Drobek M. 2022. One fold, many functions-M23 family of peptidoglycan hydrolases. Front Microbiol 13:1036964.

90. Yang DC, Blair KM, Taylor JA, Petersen TW, Sessler T, Tull CM, Leverich CK, Collar AL, Wyckoff TJ, Biboy J, Vollmer W, Salama NR. 2019. A genome-wide *Helicobacter pylori* morphology screen uncovers a membrane-spanning helical cell shape complex. J Bacteriol 201:10.1128/jb.00724-18.

91. Sichel SR, Bratton BP, Salama NR. 2022. Distinct regions of *H. pylori’*s bactofilin CcmA regulate protein-protein interactions to control helical cell shape. Elife 11:e80111.

92. Taylor JA, Bratton BP, Sichel SR, Blair KM, Jacobs HM, DeMeester KE, Kuru E, Gray J, Biboy J, VanNieuwenhze MS, Vollmer W, Grimes CL, Shaevitz JW, Salama NR. 2020. Distinct cytoskeletal proteins define zones of enhanced cell wall synthesis in *Helicobacter pylori*. Elife 9:e52482.

93. Jutras BL, Scott M, Parry B, Biboy J, Gray J, Vollmer W, Jacobs-Wagner C. 2016. Lyme disease and relapsing fever *Borrelia* elongate through zones of peptidoglycan synthesis that mark division sites of daughter cells. Proc Natl Acad Sci U S A 113:9162–70.

94. Mccausland JW, Kloos ZA, Irnov I, Sonnert ND, Zhou J, Putnick R, Mueller EA, Steere AC, Palm NW, Grimes CL, Jacobs-Wagner C. 2025. Bacterial and host enzymes modulate the inflammatory response produced by the peptidoglycan of the Lyme disease agent. BioRxiv doi:10.1101/2025.01.08.631998.

95. Takacs CN, Scott M, Chang Y, Kloos ZA, Irnov I, Rosa PA, Liu J, Jacobs-Wagner C. 2021. A CRISPR interference platform for selective downregulation of gene expression in *Borrelia burgdorferi*. Appl Environ Microbiol 87:e02519–20.

96. Di L, Pagan PE, Packer D, Martin CL, Akther S, Ramrattan G, Mongodin EF, Fraser CM, Schutzer SE, Luft BJ, Casjens SR, Qiu WG. 2014. BorreliaBase: a phylogeny-centered browser of *Borrelia* genomes. BMC Bioinformatics 15:233.

97. Abramson J, Adler J, Dunger J, Evans R, Green T, Pritzel A, Ronneberger O, Willmore L, Ballard AJ, Bambrick J, Bodenstein SW, Evans DA, Hung CC, O’Neill M, Reiman D, Tunyasuvunakool K, Wu Z, Zemgulyte A, Arvaniti E, Beattie C, Bertolli O, Bridgland A, Cherepanov A, Congreve M, Cowen-Rivers AI, Cowie A, Figurnov M, Fuchs FB, Gladman H, Jain R, Khan YA, Low CMR, Perlin K, Potapenko A, Savy P, Singh S, Stecula A, Thillaisundaram A, Tong C, Yakneen S, Zhong ED, Zielinski M, Zidek A, Bapst V, Kohli P, Jaderberg M, Hassabis D, Jumper JM. 2024. Accurate structure prediction of biomolecular interactions with AlphaFold 3. Nature 630:493–500.

98. Vasa S, Lin L, Shi C, Habenstein B, Riedel D, Kuhn J, Thanbichler M, Lange A. 2015. beta-Helical architecture of cytoskeletal bactofilin filaments revealed by solid-state NMR. Proc Natl Acad Sci U S A 112:E127–36.

99. Adams PP, Flores Avile C, Popitsch N, Bilusic I, Schroeder R, Lybecker M, Jewett MW. 2017. *In vivo* expression technology and 5’ end mapping of the *Borrelia burgdorferi* transcriptome identify novel RNAs expressed during mammalian infection. Nucleic Acids Res 45:775–792.

100. Gilbert MA, Morton EA, Bundle SF, Samuels DS. 2007. Artificial regulation of *ospC* expression in *Borrelia burgdorferi*. Mol Microbiol 63:1259–73.

101. Blevins JS, Revel AT, Smith AH, Bachlani GN, Norgard MV. 2007. Adaptation of a luciferase gene reporter and *lac* expression system to *Borrelia burgdorferi*. Appl Environ Microbiol 73:1501–13.

102. Chu CY, Stewart PE, Bestor A, Hansen B, Lin T, Gao L, Norris SJ, Rosa PA. 2016. Function of the *Borrelia burgdorferi* FtsH homolog is essential for viability both in vitro and in vivo and independent of HflK/C. mBio 7:e00404–16.

103. Ge Y, Old IG, Saint Girons I, Charon NW. 1997. Molecular characterization of a large *Borrelia burgdorferi* motility operon which is initiated by a consensus sigma70 promoter. J Bacteriol 179:2289–99.

104. Hay NA, Tipper DJ, Gygi D, Hughes C. 1999. A novel membrane protein influencing cell shape and multicellular swarming of *Proteus mirabilis*. J Bacteriol 181:2008–16.

105. Koch MK, McHugh CA, Hoiczyk E. 2011. BacM, an N-terminally processed bactofilin of *Myxococcus xanthus*, is crucial for proper cell shape. Mol Microbiol 80:1031–51.

106. Kuru E, Hughes HV, Brown PJ, Hall E, Tekkam S, Cava F, de Pedro MA, Brun YV, VanNieuwenhze MS. 2012. In situ probing of newly synthesized peptidoglycan in live bacteria with fluorescent D-amino acids. Angew Chem Int Ed Engl 51:12519–23.

107. El Andari J, Altegoer F, Bange G, Graumann PL. 2015. *Bacillus subtilis* bactofilins are essential for flagellar hook- and filament assembly and dynamically localize into structures of less than 100 nm diameter underneath the cell membrane. PLoS One 10:e0141546.

108. Kemege KE, Hickey JM, Barta ML, Wickstrum J, Balwalli N, Lovell S, Battaile KP, Hefty PS. 2015. *Chlamydia trachomatis* protein CT009 is a structural and functional homolog to the key morphogenesis component RodZ and interacts with division septal plane localized MreB. Mol Microbiol 95:365–82.

109. Ouellette SP, Karimova G, Subtil A, Ladant D. 2012. *Chlamydia* co-opts the rod shape-determining proteins MreB and Pbp2 for cell division. Mol Microbiol 85:164–78.

110. Lee J, Cox JV, Ouellette SP. 2020. Critical role for the extended N terminus of Chlamydial MreB in directing its membrane association and potential interaction with divisome proteins. J Bacteriol 202:10.1128/jb.00034-20.

111. Ouellette SP, Lee J, Cox JV. 2020. Division without binary fission: Cell division in the FtsZ-less *Chlamydia*. J Bacteriol 202:10.1128/jb.00252-20.

112. Fischbach MA, Walsh CT, Clardy J. 2008. The evolution of gene collectives: How natural selection drives chemical innovation. Proc Natl Acad Sci U S A 105:4601–8.

113. Barbour AG. 1984. Isolation and cultivation of Lyme disease spirochetes. Yale J Biol Med 57:521–5.

114. Zuckert WR. 2007. Laboratory maintenance of *Borrelia burgdorferi*. Curr Protoc Microbiol Chapter 12:Unit 12C 1.

115. Jutras BL, Chenail AM, Stevenson B. 2013. Changes in bacterial growth rate govern expression of the *Borrelia burgdorferi* OspC and Erp infection-associated surface proteins. J Bacteriol 195:757–64.

116. Takacs CN, Kloos ZA, Scott M, Rosa PA, Jacobs-Wagner C. 2018. Fluorescent proteins, promoters, and selectable markers for applications in the Lyme disease spirochete *Borrelia burgdorferi*. Appl Environ Microbiol 84:e01824–18.

117. Frank KL, Bundle SF, Kresge ME, Eggers CH, Samuels DS. 2003. *aadA* confers streptomycin resistance in *Borrelia burgdorferi*. J Bacteriol 185:6723–7.

118. Elias AF, Bono JL, Kupko JJ, 3rd, Stewart PE, Krum JG, Rosa PA. 2003. New antibiotic resistance cassettes suitable for genetic studies in *Borrelia burgdorferi*. J Mol Microbiol Biotechnol 6:29–40.

119. Bono JL, Elias AF, Kupko JJ, 3rd, Stevenson B, Tilly K, Rosa P. 2000. Efficient targeted mutagenesis in *Borrelia burgdorferi*. J Bacteriol 182:2445–52.

120. Takacs CN, Wachter J, Xiang Y, Ren Z, Karaboja X, Scott M, Stoner MR, Irnov I, Jannetty N, Rosa PA, Wang X, Jacobs-Wagner C. 2022. Polyploidy, regular patterning of genome copies, and unusual control of DNA partitioning in the Lyme disease spirochete. Nat Commun 13:7173.

121. Tilly K, Elias AF, Bono JL, Stewart P, Rosa P. 2000. DNA exchange and insertional inactivation in spirochetes. J Mol Microbiol Biotechnol 2:433–42.

122. Samuels DS, Drecktrah D, Hall LS. 2018. Genetic transformation and complementation. Methods Mol Biol 1690:183–200.

123. Samuels DS. 1995. Electrotransformation of the spirochete *Borrelia burgdorferi*. Methods Mol Biol 47:253–9.

124. Green MR, Sambrook J. 2016. Precipitation of DNA with ethanol. Cold Spring Harb Protoc 2016:pdb.prot093377.

125. Bunikis I, Kutschan-Bunikis S, Bonde M, Bergstrom S. 2011. Multiplex PCR as a tool for validating plasmid content of *Borrelia burgdorferi*. J Microbiol Methods 86:243–7.

126. Nowalk AJ, Gilmore RD, Jr., Carroll JA. 2006. Serologic proteome analysis of *Borrelia burgdorferi* membrane-associated proteins. Infect Immun 74:3864–73.

127. Livak KJ, Schmittgen TD. 2001. Analysis of relative gene expression data using real-time quantitative PCR and the 2(-Delta Delta C(T)) Method. Methods 25:402–8.

128. Wachter J, Cheff B, Hillman C, Carracoi V, Dorward DW, Martens C, Barbian K, Nardone G, Renee Olano L, Kinnersley M, Secor PR, Rosa PA. 2023. Coupled induction of prophage and virulence factors during tick transmission of the Lyme disease spirochete. Nat Commun 14:198.

129. Langmead B, Salzberg SL. 2012. Fast gapped-read alignment with Bowtie 2. Nat Methods 9:357–9.

130. Liao Y, Smyth GK, Shi W. 2019. The R package Rsubread is easier, faster, cheaper and better for alignment and quantification of RNA sequencing reads. Nucleic Acids Res 47:e47.

131. Liao Y, Smyth GK, Shi W. 2014. featureCounts: an efficient general purpose program for assigning sequence reads to genomic features. Bioinformatics 30:923–30.

132. Glaser P, Sharpe ME, Raether B, Perego M, Ohlsen K, Errington J. 1997. Dynamic, mitotic-like behavior of a bacterial protein required for accurate chromosome partitioning. Genes Dev 11:1160–8.

133. Paintdakhi A, Parry B, Campos M, Irnov I, Elf J, Surovtsev I, Jacobs-Wagner C. 2016. Oufti: an integrated software package for high-accuracy, high-throughput quantitative microscopy analysis. Mol Microbiol 99:767–77.

134. Schindelin J, Arganda-Carreras I, Frise E, Kaynig V, Longair M, Pietzsch T, Preibisch S, Rueden C, Saalfeld S, Schmid B, Tinevez JY, White DJ, Hartenstein V, Eliceiri K, Tomancak P, Cardona A. 2012. Fiji: an open-source platform for biological-image analysis. Nat Methods 9:676–82.

135. Camacho C, Coulouris G, Avagyan V, Ma N, Papadopoulos J, Bealer K, Madden TL. 2009. BLAST+: architecture and applications. BMC Bioinformatics 10:421.

136. Jumper J, Evans R, Pritzel A, Green T, Figurnov M, Ronneberger O, Tunyasuvunakool K, Bates R, Zidek A, Potapenko A, Bridgland A, Meyer C, Kohl SAA, Ballard AJ, Cowie A, Romera-Paredes B, Nikolov S, Jain R, Adler J, Back T, Petersen S, Reiman D, Clancy E, Zielinski M, Steinegger M, Pacholska M, Berghammer T, Bodenstein S, Silver D, Vinyals O, Senior AW, Kavukcuoglu K, Kohli P, Hassabis D. 2021. Highly accurate protein structure prediction with AlphaFold. Nature 596:583–589.

137. Mirdita M, Schutze K, Moriwaki Y, Heo L, Ovchinnikov S, Steinegger M. 2022. ColabFold: making protein folding accessible to all. Nat Methods 19:679–682.

138. Pettersen EF, Goddard TD, Huang CC, Meng EC, Couch GS, Croll TI, Morris JH, Ferrin TE. 2021. UCSF ChimeraX: Structure visualization for researchers, educators, and developers. Protein Sci 30:70–82.

139. Fraser CM, Casjens S, Huang WM, Sutton GG, Clayton R, Lathigra R, White O, Ketchum KA, Dodson R, Hickey EK, Gwinn M, Dougherty B, Tomb JF, Fleischmann RD, Richardson D, Peterson J, Kerlavage AR, Quackenbush J, Salzberg S, Hanson M, van Vugt R, Palmer N, Adams MD, Gocayne J, Weidman J, Utterback T, Watthey L, McDonald L, Artiach P, Bowman C, Garland S, Fuji C, Cotton MD, Horst K, Roberts K, Hatch B, Smith HO, Venter JC. 1997. Genomic sequence of a Lyme disease spirochaete, *Borrelia burgdorferi*. Nature 390:580–6.

140. Gross LA, Baird GS, Hoffman RC, Baldridge KK, Tsien RY. 2000. The structure of the chromophore within DsRed, a red fluorescent protein from coral. Proc Natl Acad Sci U S A 97:11990–5.

141. Rego RO, Bestor A, Rosa PA. 2011. Defining the plasmid-borne restriction-modification systems of the Lyme disease spirochete *Borrelia burgdorferi*. J Bacteriol 193:1161–71.

142. Stewart PE, Thalken R, Bono JL, Rosa P. 2001. Isolation of a circular plasmid region sufficient for autonomous replication and transformation of infectious *Borrelia burgdorferi*. Mol Microbiol 39:714–21.

143. Narasimhan S, Sukumaran B, Bozdogan U, Thomas V, Liang X, DePonte K, Marcantonio N, Koski RA, Anderson JF, Kantor F, Fikrig E. 2007. A tick antioxidant facilitates the Lyme disease agent’s successful migration from the mammalian host to the arthropod vector. Cell Host Microbe 2:7–18.

